# Natural enemies can mediate the impact of plant microbiota on insect-borne virus transmission

**DOI:** 10.64898/2026.05.11.724421

**Authors:** Lucas A. Nell, Tory A. Hendry, Andrew M. Hein, Megan A. Greischar

## Abstract

Individual hosts can be protected from disease vectors through their microbiota, but how those benefits scale up to the host population level is rarely obvious. Disease spread may be inhibited through reduced vector visitation or amplified as vectors redirect their attention to unprotected hosts. The outcome—suppression or escalation of disease outbreaks—is likely to be influenced by natural enemies that redistribute or suppress vector populations. The challenge of determining how these processes interact across scales is exemplified by pea aphids, major virus vectors in pea crops that are commonly managed using parasitoid wasps. Recent evidence suggests that epiphytic bacteria in the genus *Pseudomonas* can repel or kill pea aphids, yet whether *Pseudomonas*-mediated protection from aphids complements or undermines parasitoid-based vector control remains unknown. We developed a multi-scale mathematical model to investigate why *Pseudomonas* complements versus undermines biocontrol of aphid-vectored virus outbreaks. Because *Pseudomonas*-induced mortality leaves aphids patchily distributed, the effect of *Pseudomonas* on virus outbreaks depends most strongly on how well parasitoids track aphid densities: When parasitoids efficiently locate aphids, *Pseudomonas* inhibits virus outbreaks by reducing aphid densities. However, when parasitoids are less efficient, they often search unproductively for aphids on *Pseudomonas*-inhabited plants with few aphids, slowing parasitoid population growth. Reduced parasitism intensifies aphid crowding on *Pseudomonas*-free plants and boosts the production of winged aphids that accelerate viral spread. Counterintuitively, the more effective *Pseudomonas* is at killing aphids, the more strongly it generates spatial variability and promotes virus spread. Our results show how community context dictates whether individual microbiota protection slows or accelerates viral outbreaks at the population scale.

## Introduction

Microbiomes vary across hosts and influence attractiveness to disease vectors (reviewed in Grunseich et al. 2019), leaving a subset of hosts protected by virtue of being less attractive to vectors. While protection of some hosts could limit disease spread (e.g., through herd immunity), it could also divert vectors to unprotected individuals (as observed with mosquito repellent, Moore et al. 2007). Theory predicts that when vectors are diverted from protected hosts to unprotected hosts that are more efficient at transmitting the pathogen, partial population protection can accelerate disease spread beyond what would occur if no hosts were protected at all (Miller et al. 2016). The risks of partial protection could be exacerbated by two aspects community ecology that warrant further attention: First, availability of preferred hosts impacts vector population dynamics and hence potential for disease spread. Second, natural enemies of vectors can curb disease risk (Wyckhuys et al. 2025), but their ability to find and remove vectors depends on how vectors are distributed across hosts. Thus, variation in microbiomes could amplify or curtail disease spread, depending on interactions between vector population dynamics and natural enemies.

Vector feeding preferences have been extensively studied in plants, where a range of cues inform the movement decisions of individual vectors (reviewed in Bruce et al. 2005, Eigenbrode et al. 2018). Plant microbiota mediate many of the cues used by insect vectors (reviewed in Zhang et al. 2024, Grunseich et al. 2019), including through changes in plant-produced volatile organic compounds (VOCs) (Bruce et al. 2005, Rostás et al. 2015). Microbiota on the surface of plants can also include entomopathogens that are visually detected and avoided by insect herbivores, such as spongy moth (*Lymantria dispar*) larvae that can avoid conspecific cadavers infected by baculoviruses (Capinera et al. 1976) or aphid vectors that avoid entomopathogenic members of plant surface microbiomes (Hendry et al. 2018). Distinct from symbiotic or commensal components of the plant microbiota, plants infected with phytopathogens can be more attractive to insect vectors (reviewed in Eigenbrode et al. 2018), which can hasten disease spread (Shaw et al. 2017, Shoemaker et al. 2019).

Natural enemies like parasitoids have the potential to reduce vector-borne disease transmission. A meta-analysis of field studies suggests that vector-borne pathogen incidence scales with vector abundance and that natural enemies usually reduce vector abundance (Wyckhuys et al. 2025). Yet theory suggests potential for complex links between natural enemies, vector abundance, and vector-borne disease prevalence, including the possibility of parasitoid suppression of outbreaks despite limited impact on vector abundance (Okamoto and Amarasekare 2012). A key driver of parasitoid persistence—and hence potential for sustained biocontrol—is parasitoid efficiency in finding hosts and aggregation of host and parasitoid populations (Hassell and May 1973, reviewed in Godfray and Pacala 1992). Aggregation of the vectors targeted by parasitoids depends in part on vector dispersal, which is a function of population density for many insect vectors (Braendle et al. 2006, Roff 1986, Xu and Zhang 2017, Bosch and Ives 2023). For these insects, dispersal morphotypes are produced based on developmental signals from mothers to offspring before birth (Sutherland 1969, Denno 1994). The production of these signals is thought to be affected by a number of cues, but local population density is one of the most consistent (Bosch and Ives 2023, Denno 1994, Braendle et al. 2006). Dispersal morphotypes are better suited to escape poor local conditions and probe new plants until settling to feed indefinitely, so their production is predicted to be especially important in determining how these insect vectors spread pathogens (Donnelly et al. 2019).

Pea aphids (*Acyrthosiphon pisum*) represent important vectors of plant viruses and a data-rich system in which to build understanding for how microbiota-mediated protection of individual plants impacts epidemiological dynamics. Pea aphids are a common sap-sucking insect pest of legumes such as pea (*Pisum sativum*). Aphids mostly impact pea plants by vectoring viruses (e.g., *Potyvirus* sp., *Enamovirus* sp., Ng and Perry 2004). As aphid density increases on plants, an increasing fraction of offspring develop into winged forms called “alates” (Sutherland 1969, Bosch and Ives 2023, Nell et al. 2024), a distinct morphotype that enables spread of virus to new plants. In addition to vectoring plant viruses, pea aphids also contend with their own pathogens. *Pseudomonas* is a genus of bacteria that are commonly found on plant leaf surfaces, and some strains can be a highly virulent pathogen of pea aphids (Hendry et al. 2014). Pea aphids can detect the ultraviolet-based fluorescence of some virulent strains of *P. syringae* and avoid them (Hendry et al. 2018). The presence of entomopathogenic, fluorescent strains of *Pseudomonas* (hereafter simply *Pseudomonas*) on individual pea plants should ostensibly reduce virus transmission. However, *Pseudomonas* repelling aphids may cause crowding in non- or low-*Pseudomonas* plants, which could result in greater abundances of alates across plant populations. Given that alates disperse more readily than non-winged forms, *Pseudomonas* could promote virus outbreaks. In addition, patchy *Pseudomonas* distributions and *Pseudomonas*-induced aphid mortality could generate spatial variability in aphid densities that reduces natural enemy efficiency. Biocontrol of aphid populations has long been of interest in this system, including the parasitoid wasp *Aphidius ervi*, which can greatly suppress aphid populations (Ives et al. 1999, Nell et al. 2024). However, differences in aphid abundance across plants reduce parasitoid searching efficiency (Ives et al. 1999). Thus, *Pseudomonas*-induced heterogeneity in aphid abundance across plants has the potential to undermine biological control of *A. ervi* parasitoids and drive viral outbreaks (Figure 1A).

**Figure 1:**
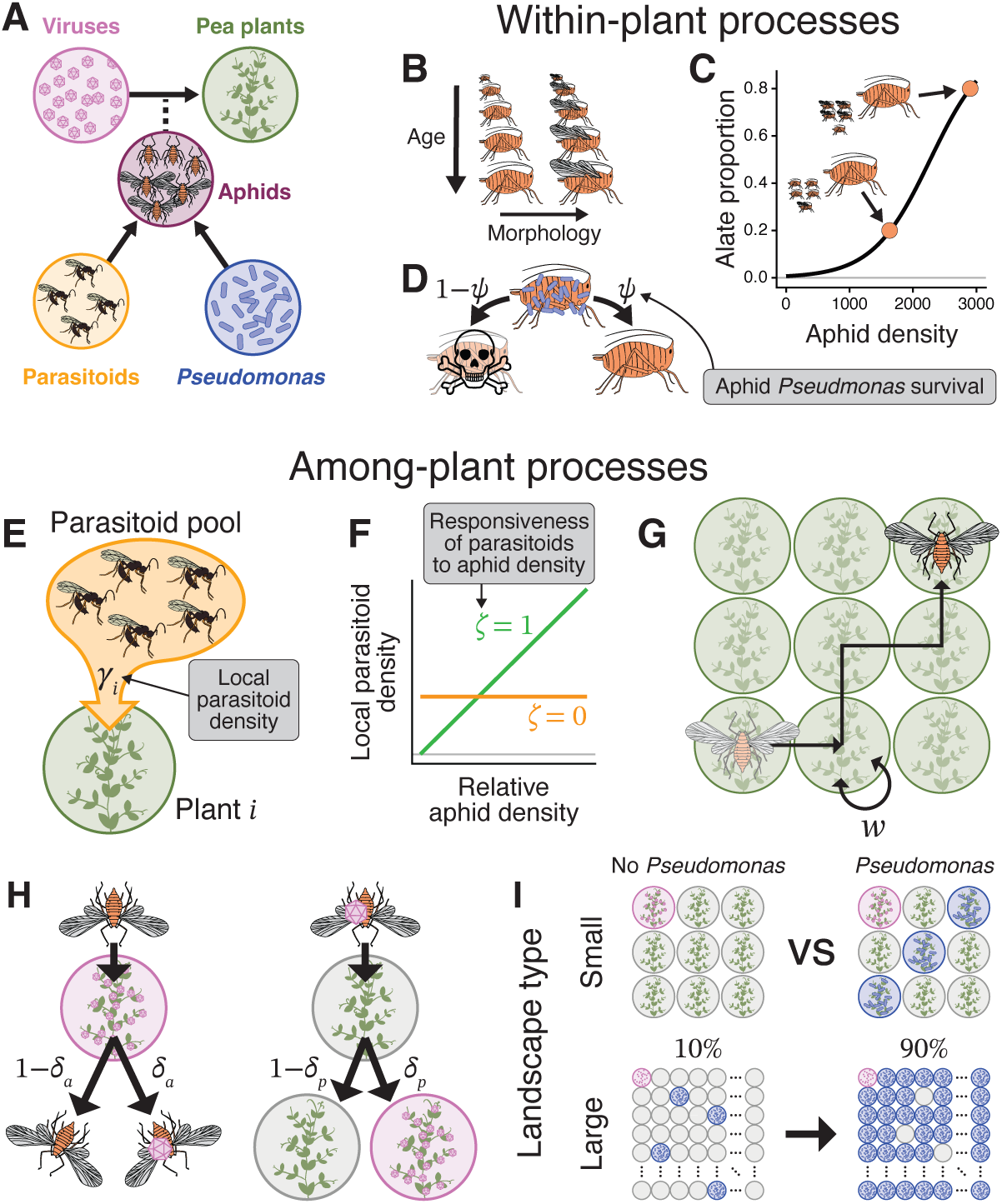
The spread of aphid-borne viruses depends on dynamics at multiple scales, incorporated into a multi-scale model of viral spread through a field of pea plants. (A) We model key species interactions, tracking how aphids vector plant viruses and the impact of parasitoids and *Pseudomonas* on aphid populations. (B) Aphid populations are structured by age and by morphology (winged “alates” and non-winged). (C) Alate production is dictated by total aphids (alates, non-winged, and parasitized) on the plant. (D) Aphids have a 1 − *ψ* chance per day of being killed by *Pseudomonas* on plants with the bacteria. (E) Densities of adult, female parasitoids at individual plants (*γ*) are drawn from the landscape-wide parasitoid pool. (F) Parasitoid responsiveness to aphid densities (*ζ*) shapes the functional form of the *γ*–aphid density relationship. (G) Dispersing alates probe successive plants until they settle to feed with probability *w*. (H) Along the flight path, alates acquire virus from plants and inoculate plants with virus with probabilities *δ_a_* and *δ_p_*, respectively. (I) We simulated two landscape types: Small landscapes contain zero or three *Pseudomonas*-inhabited plants (blue). Large landscapes contain zero (not shown) or 10–90% *Pseudomonas*-inhabited plants. Virus starts in the upper left corner (pink). Illustrations by Dr. Esther Nell; used with permission.

Here, we use a mathematical model of pea aphid–parasitoid interactions and aphid-borne virus transmission to understand when and why individual-plant protection by *Pseudomonas* promotes or inhibits viral outbreaks at the population scale. We find that *Pseudomonas* offers population-level protection—reducing virus transmission—when parasitoids efficiently locate aphid populations. However, *Pseudomonas* can promote virus transmission when parasitoid searching efficiency is low, since parasitoids waste time searching *Pseudomonas*-protected plants with few aphids. As a result, parasitoid numbers are slower to increase in response to aphid abundance, and aphids reach high densities on unprotected plants, promoting alate production and subsequent viral spread. This counter-productive outcome is exacerbated with reductions in aphid survival on *Pseudomonas*-protected plants because that amplifies the variation in aphid densities across plants, rendering parasitoids even less efficient at locating and controlling aphid populations. Stronger parasitoid responsiveness to aphid densities always pushes *Pseudomonas* towards a more inhibitory effect on virus transmission. Our results show that vector–natural enemy interactions can determine how individual host protection affects population-level disease transmission.

## Methods

To understanding the factors that change how *Pseudomonas* affects vector-borne disease outbreaks, we used a mathematical model that explicitly simulates both within- and among-plant processes for a single field of pea plants. Within-plant processes include stage-structured aphid population growth (Figure 1B), production of winged alates (Figure 1C), and *Pseudomonas*-induced aphid mortality (Figure 1D). We described these processes with a stage-structured, pea aphid–parasitoid model based on previous studies (Ives et al. 2020, Meisner et al. 2014, Nell et al. 2024). Among-plant processes include adult parasitoid densities (Figure 1E,F), alate dispersal (Figure 1G), virus spread (Figure 1H), and the density and distribution of *Pseudomonas* (Figure 1I). We assume that *Pseudomonas* does not move across the landscape, due to the unknowns of *Pseudomonas* dispersal (detailed in section “Spatial and disease model” below). We used a spatially explicit, individual-based model of alate dispersal and vector-mediated, nonpersistent virus transmission based on Donnelly et al. (2019). Combining these approaches allowed us to use previous work describing pea aphid–parasitoid dynamics at local scales, while incorporating spatial structure in our simulation of virus transmission.

We first simulated small (3 × 3 grid of individual plants) landscapes to understand how population and community processes affect virus transmission when other factors (e.g., *Pseudomonas* repellence, varying spatial configurations) are not at play. We then simulated larger (100 × 100 grid) landscapes to understand how factors related to spatial variability affect virus transmission and modulate the impact of *Pseudomonas*. We lastly compared simulations to previously reported experimental microcosm data (Ives et al. 1999) to estimate parasitoid responsiveness to aphid densities.

### Host–parasitoid model

We started with an aphid–parasitoid model that has been shown to accurately capture the dynamics of this system for both the field and laboratory (Ives et al. 2020, Meisner et al. 2014, Nell et al. 2024). This model has been previously described in detail (Nell et al. 2024), so the model equations from previous versions are in Appendix S1: Section S1. The main differences from previous versions are that here, we (i) simulate multiple plants within a single field, (ii) assume a monoclonal aphid population (to exclude the effects of evolution), and (iii) include aphid infection by *Pseudomonas*. Adult parasitoids are far more mobile than aphids, so this single-field version tracks adult, female parasitoids as one field-wide pool, *Y*. To estimate aphid survival from parasitoid attack, we used a new equation for the density of parasitoids searching for aphid hosts on plant *i* (*γ*(·)):

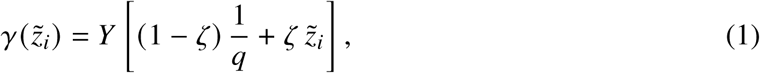

where *z*°*_i_* is the relative number of aphids on plant *i* (i.e., total aphids on plant *i* divided by total aphids on landscape), *q* is the number of plants, and *ζ* ∈ [0, 1] governs the functional form of the relationship between *γ* and aphid density (Godfray and Pacala 1992). When *ζ* = 0, parasitoid densities are evenly distributed regardless of aphid densities, whereas when *ζ* = 1, there is a one-to-one relationship between relative parasitoid and aphid densities.

### Spatial and disease model

Landscapes represent a single field of *q* total pea plants. We used a single field because alates typically notice *Pseudomonas* only once they are close to the plant (≤ 180 cm), and *Pseudomonas* densities can vary significantly even at fine spatial scales (∼1–5 meters) (Herr et al. 2024). We model aphid populations at the individual-plant level rather than as a single field-wide population because alate production is driven primarily by aphid density on a particular plant (Bosch and Ives 2023). *Pseudomonas* is either present (*ψ* < 1) or absent (*ψ* = 1) on each plant, and the distribution of *Pseudomonas* does not change over the course of each simulation, since little is known about how *Pseudomonas* disperses across the landscape and whether aphids (through infection or by consuming honeydew) affect this process. One plant starts as infectious with the virus, and the virus is only transmitted between plants through the dispersal of alates. We assumed that all virus transmission is via nonpersistent transmission, where the virus transiently contaminates alate mouthparts, transmission biology shared by most aphid-borne viruses (Nault 1997). Thus, we assume alates can only transmit viruses between plants visited on the same day. We assumed that infectious plants do not recover, a characteristic of many aphid-vectored viruses (e.g., pea seedborne mosaic viruses, Roberts et al. 2003).

#### Alate dispersal

Alates are the only aphids that move between plants or vector diseases in our model. Every day, the number of adult alates of age *k* that leaves their starting plant (*n*_fly,*k*_) is sampled as ∼ Binomial (⌊*X_k_* ⌋, *p*_fly_), where ⌊·⌋ is the floor function (to coerce to an integer without potentially adding aphids), *X_k_* is the density of adult alates of stage *k*, and *p*_fly_ is the probability that an adult alate leaves their plant each day. Alate flight paths are simulated individually. Each alate that leaves a plant can travel to any other plant that is within alates’ dispersal radius (*r*_fly_). Once an alate flies to a new plant, it probes the plant, and will stay on that plant with probability *w*. It will continue to visit and probe plants until it chooses to stay indefinitely. We assume that alates visiting plants with *Pseudomonas* then leaving the same day are not exposed for a long enough period to be infected by the bacteria. In the main text, we show simulations where plants are equally attractive to alates, but we also simulated scenarios where alates were attracted to virus-infected plants and repelled from *Pseudomonas*-inhabited plants. These factors could change virus transmission but did not substantially alter the relationship between *Pseudomonas* and virus outbreaks (Appendix S1: Section S2).

#### Virus spread

We assume that alates probe plants on each visit, which is required for virus transmission (Krenz et al. 2015, Powell 2005). If an alate visits a plant infectious with the virus (including the plant they start on), the virus loads onto that alate with probability *δ_a_*. If a virus-loaded alate probes an uninfected plant, the probability that the plant is exposed to the virus is *δ_p_*. A plant that is exposed to the virus is not immediately able to pass the virus onto other alates. Plants transition from exposed to infectious after 7 days, the approximate length of time required for cucumber mosaic virus (Davis and Hampton 1986), which spreads via nonpersistent transmission by aphids. We assumed that once an alate is loaded with the virus, it can deliver the virus to all subsequent plants it probes on that day. After that day, alates lose the virus unless they settle on an infectious plant, in which case they acquire it with probability *δ_a_*.

### Parameterization and simulation configurations

In all simulations, we simulated 100 days, a typical growing season for pea plant crops, and the plant in the upper-left corner of the landscape (*x* = 1, *y* = 1) always started as infectious with the virus. Although this approach may produce edge effects, we expect that initially infectious plants in the field would be biased toward plants near edges. Moreover, simulating interior starting locations did not change our conclusions (Appendix S1: Section S3). We also started every plant with the same total density of non-winged aphids (*N*_0_). This total density was split among age stages by assuming aphids were at their stable age distribution. We varied starting aphid and parasitoid densities in small landscapes, as well as aphid carrying capacities, but these factors only exaggerated effects of *Pseudomonas* as dictated by other parameters (Appendix S1: Section S4). Similarly, we also simulated variable starting aphid densities in large landscapes, but this had only small effects (Appendix S1: Section S5). In all simulations, plants start with no alates, parasitized aphids, or mummies.

To first understand how population and community processes interact to change disease dynamics, we conducted simulations of small 3 × 3 landscapes, where *Pseudomonas* always started on the back diagonal of the landscape (i.e, (*x*, *y*) = (1, 3), (2, 2), and (3, 1)) (Figure 1I). This configuration produced similar mean peak infected plants across 1,000 simulations to when *Pseudomonas* was randomly located on the landscape, but with reduced among-simulation variation. This smaller size allowed us to more efficiently simulate across a wider range of possible parameter values before moving on to larger landscapes.

To next explore factors related to spatial variability, we used 100 × 100 landscapes that more closely match the size of a typical agricultural field (Figure 1I). Assuming spacing of 0.75 meters (typical for fresh pea), each landscape is 0.5625 hectare. Within these landscapes, we simulated *Pseudomonas* occupying 0, 1000, 3000, 5000, 7000, or 9000 plants. This allowed us to compare across a wide range of *Pseudomonas* occupancies spanning the 59% pea plant leaves inhabited by fluorescent *Pseudomonas* estimated by Herr et al. (2024). *Pseudomonas* locations in these landscapes were spatially uniform (i.e., all plants had an equal probability of being occupied by *Pseudomonas*). Clustered *Pseudomonas* locations had no effect on virus transmission (Appendix S1: Section S3). The initially infectious plant was not inhabited by *Pseudomonas*. The only effect of starting the virus on a *Pseudomonas*-inhabited plant is suppressed virus transmission, driven by reduced alate production on plants where aphids are killed by *Pseudomonas* (Appendix S1: Section S6).

In the main text, we use peak infected plants as our transmission metric; because recovery does not occur, this value also represents the final count. This metric effectively captures transmission trends. For completeness, however, we provide supplementary figures for scenarios where outbreaks (i.e., at least one new viral infection) do not always occur, reporting both the probability of emergence (i.e., an outbreak occurring) and the mean outbreak size conditional on emergence. We also report large-landscape simulations with very small loading probabilities (*δ_a_* = *δ_p_* = 0.05) which caused outbreaks to be comparatively rare; our results were unchanged (Appendix S1: Section S3). Because simulations are stochastic, we also show 95% confidence intervals for means of all virus-infection outcomes, calculated using nonparametric bootstrapping. See Appendix S1: Table S1 for state variables, starting conditions, and parameter values, and Appendix S1: Section S7 for model parameterization.

### Estimating parasitoid responsiveness (ζ)

We lastly conducted simulations aimed at estimating a reasonable value for parasitoid responsiveness to aphid densities (*ζ*). Ives et al. (1999) estimated foraging times for parasitoid *A. ervi* as a function of pea aphid densities. They found that parasitoids spent about 3.76× more time on a plant if they encountered at least one aphid, and the probability of an encounter increased with aphid density. Aphid densities in their small experimental microcosms (75 × 150 × 5 cm, containing 32 plants) are not directly applicable to plants in a field, which precluded us from using aphid densities to directly calculate parasitoid densities based on their results. Instead, we assumed that parasitoids always encounter aphids on *Pseudomonas*-free plants and never do in *Pseudomonas*-inhabited plants when aphids are at peak densities (and spatial variation in densities is maximized). Under this assumption, in a landscape of *q* total plants, *q_p_* of which are inhabited by *Pseudomonas*, the expected proportion of time parasitoids spend foraging on a given *Pseudomonas*-free plant is 3.76/(3.76 (*q* − *q_p_*) + *q_p_*), and 1/(3.76 (*q* − *q_p_*) + *q_p_*) for *Pseudomonas*-inhabited plants. We assumed that the expected plant-level density of parasitoids was the product of the total parasitoid density and the expected proportion of time parasitoids spend on each plant. Because this depends on the number of *Pseudomonas*-inhabited plants, we conducted simulations of large landscapes containing 1000, 3000, 5000, 7000, or 9000 *Pseudomonas*-inhabited plants across a range of *ζ* values. We then filtered each simulation for the time step when aphid density was maximized. At this time step, we compared the observed parasitoid density from the simulations to the expected density based on Ives et al. (1999) to visualize which value of *ζ* most closely matched their results across levels of *Pseudomonas* occupancy.

## Results

### *Pseudomonas* can promote viral spread by hindering parasitoids

Focusing first on small 3 × 3 plots, we find that *Pseudomonas* can promote virus spread when parasitoids respond weakly to aphid densities (*ζ* = 0.1), even when alates are not repelled by the presence of *Pseudomonas* (i.e., *ρ* = 1). In this case, *Pseudomonas* reduces peak densities of total aphids and parasitoids, but it increases peak densities of adult alates and the number of infected plants (Figure 2A). This occurs because *Pseudomonas*-induced aphid mortality on some, but not all, plants generates spatial variability in aphid densities. When combined with poor tracking of aphids by parasitoids, variability in aphid densities causes some adult parasitoids to spend time searching for aphids where there are effectively none (Figure 2C). Inefficiency in parasitoid attacks slows parasitoid population growth (Figure 2A) and reduces parasitoid densities on *Pseudomonas*-free plants in landscapes with *Pseudomonas* (Figure 2C). For this reason, *Pseudomonas*-free plants maintain higher densities of total aphids for longer and produce more alates in *Pseudomonas*-containing landscapes compared to plants in *Pseudomonas*-free landscapes. Because of the nonlinear relationship between aphid density and alate offspring production (Figure 1C), this effect is large enough to increase total alate production in *Pseudomonas*-inhabited versus -free landscapes, despite alate production only occurring on six of nine plants (Figure 2A,C). The increase in adult alates when *Pseudomonas* is present increases the peak number of infected plants (Figure 2A,E), and the odds and size of outbreaks (Appendix S1: Figure S1A,C).

**Figure 2:**
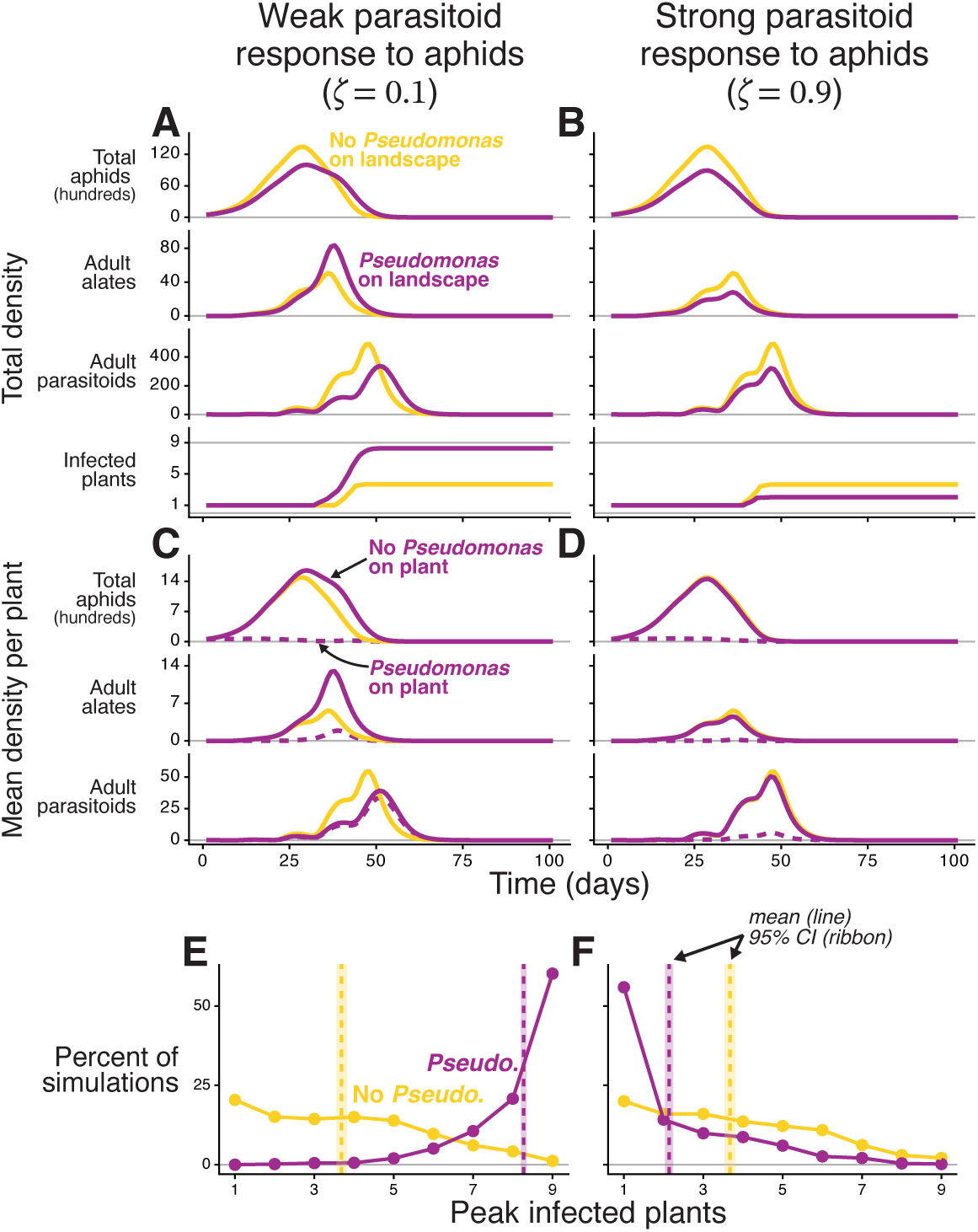
In 3 × 3 landscapes of pea plants, *Pseudomonas* can either promote or inhibit virus outbreaks depending on how strongly parasitoids respond to aphid densities. Parasitoids respond (A,C,E) weakly or (B,D,F) strongly to aphid densities. (A,B) Total aphid, adult alate, adult parasitoid (females only), and virus-infected plant densities summed across the 9 plants through time. (C,D) Mean per-plant densities through time for all but infected plants; lines for *Pseudomonas*-present landscapes are separated into whether *Pseudomonas* is absent (solid line) or or present (dashed line) on the plant. (A–D) All densities are means through time across 1000 simulations, and color indicates whether landscapes have 3 (magenta) or zero (gold) *Pseudomonas*-inhabited plants. (E,F) Frequency of peak infected plants; vertical dashed lines are means, and ribbons are bootstrapped 95% confidence intervals. For all simulations, *Y*_0_ = 1, *N*_0_ = 55, *ν* = 1, *ρ* = 1, and *ψ* = 0.85.

In contrast, when parasitoids respond strongly to aphid densities (*ζ* = 0.9), *Pseudomonas* can inhibit virus spread. Here, *Pseudomonas* presence reduces total aphid, alate, and parasitoids densities, but the timing of the peak densities remains the same regardless of *Pseudomonas* presence (Figure 2B). Moreover, per-plant densities of aphids, alates, and parasitoids are nearly identical in *Pseudomonas*-free plants regardless of *Pseudomonas* presence on the landscape (Figure 2D). When parasitoids effectively track aphids, landscapes with *Pseudomonas* only differ from those without *Pseudomonas* by having three plants where aphids are nearly extinct and alates production is negligible. This causes *Pseudomonas* presence on the landscape to decrease peak infected plants (Figure 2B,F), and the odds and size of outbreaks (Appendix S1: Figure S1B,D).

### *Pseudomonas* effects depend on parasitoid responsiveness to aphid densities

The responsiveness of parasitoids to aphid densities (*ζ*) is the main driver of whether *Pseudomonas* inhibits or promotes virus transmission (Figure 3A). Weaker parasitoid responsiveness (i.e., lower *ζ*) causes parasitoid population growth to be further slowed by the spatial variability in aphid densities generated by *Pseudomonas* presence. Slow parasitoid population growth releases aphids from parasitism, facilitating production of more virus-spreading alates. Thus, weakening parasitoid responsiveness will always increase the peak number of infected plants in *Pseudomonas*-inhabited landscapes. Parasitoid responsiveness does not affect virus transmission when *Pseudomonas* is absent from the landscape because *Pseudomonas* is the primary factor introducing spatial variability in aphid abundances. Thus weaker parasitoid responsiveness to aphids pushes *Pseudomonas* toward promoting virus transmission (Figure 3A, Appendix S1: Figure S2). For the parameter values and starting conditions used for our default small-landscape simulations, parasitoid responsiveness of *ζ* ≈ 0.7 (*ζ* = 0: parasitoids uniform; *ζ* = 1: parasitoids track aphids proportionally) is the breakpoint above which *Pseudomonas* inhibits virus transmission.

**Figure 3:**
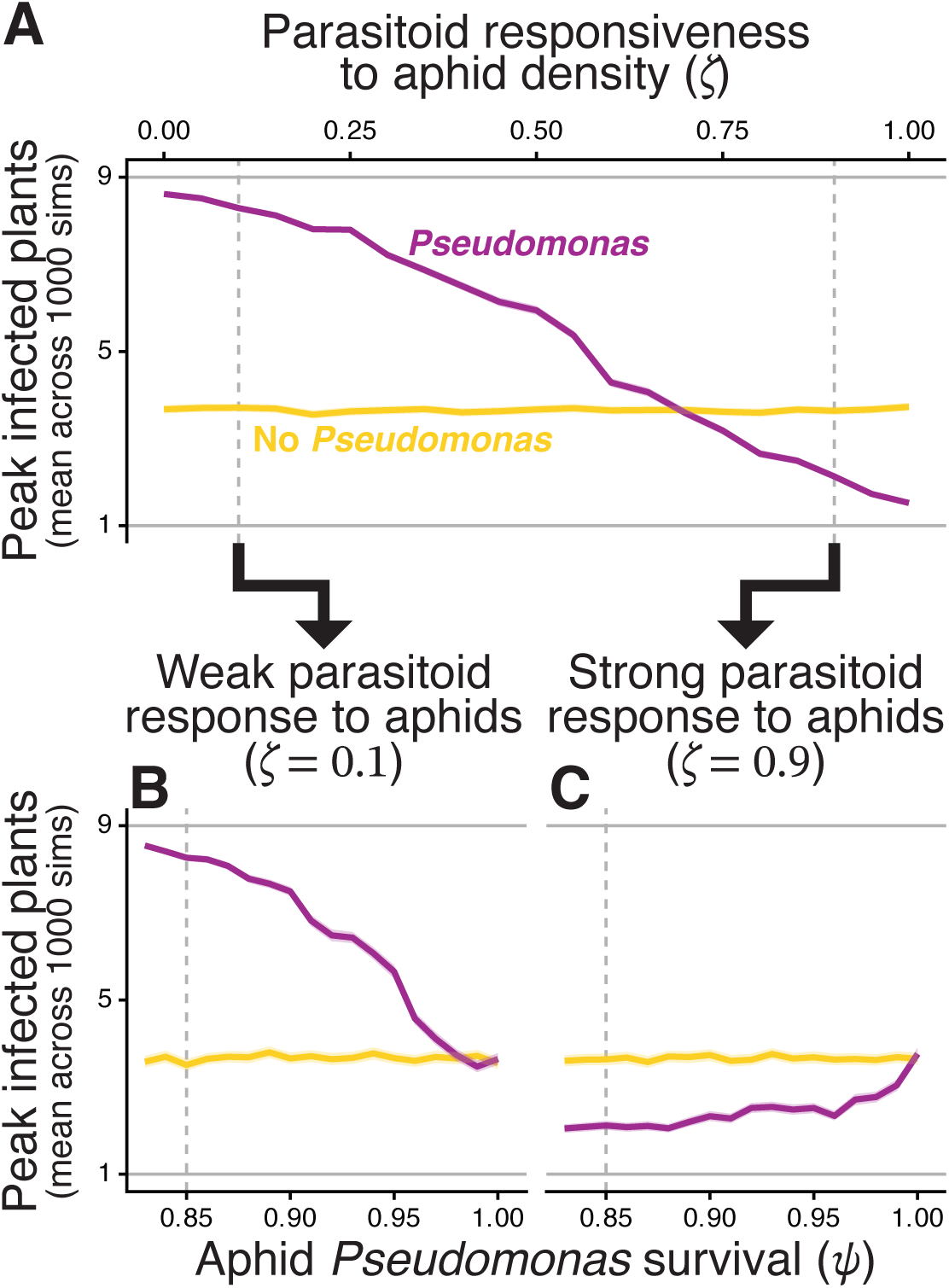
Whether *Pseudomonas* promotes or hinders virus transmission depends on parasitoid responsiveness to aphid densities (*ζ*), with the magnitude of those effects exaggerated by *Pseudomonas*-induced aphid mortality. (A) Mean peak infected plants across parasitoid responsiveness. (B,C) Mean peak infected plants versus aphid survival from *Pseudomonas* infection (*ψ*), for (B) weak or (C) strong parasitoid responsiveness. Color indicates whether landscapes have 3 (magenta) or zero (gold) *Pseudomonas*-inhabited plants; ribbons show 95% confidence intervals. Vertical dashed lines mark parameter values used in Figure 2. Except for parameters shown along the x-axis of each panel, simulation configurations match Figure 2.

To investigate the role of aphid density-dependent alate production in driving *Pseudomonas* effects, we simulated our model without an effect of aphid density on alate production and across levels of parasitoid responsiveness to aphids. When alate production is a fixed proportion of offspring, parasitoid responsiveness has a weaker effect on virus transmission, but *Pseudomonas* presence can still either inhibit or promote virus transmission (Appendix S1: Figure S3). This is because density-dependent alate production amplifies the effect of total aphid densities on alates, which strengthens the impact of partial *Pseudomonas* protection of host plants.

### Greater *Pseudomonas*-induced aphid mortality can promote viral transmission

The effect of reducing aphid *Pseudomonas* survival (*ψ*) varies depending on parasitoid responsiveness to aphid densities. When parasitoids respond weakly to aphids, decreasing aphid *Pseudomonas* survival increases the spatial variability in aphid densities generated by *Pseudomonas*, further slowing the population growth of parasitoids. This increases the number of peak infected plants for *Pseudomonas*-inhabited landscapes, strengthening the virus-promoting effects of *Pseudomonas* (Figure 3B, Appendix S1: Figure S4A,C). Thus, when parasitoids track aphid densities poorly, the more effective *Pseudomonas* is at protecting individual plants, the more it contributes to virus transmission at the population scale. In contrast, when parasitoids respond strongly to aphids, decreasing aphid *Pseudomonas* survival causes *Pseudomonas*-inhabited plants to produce fewer alates and act as stronger sinks for incoming alates from other plants, reducing alate vectors and virus spread across the landscape. This strengthens the virus-inhibiting effects of *Pseudomonas* (Figure 3C, Appendix S1: Figure S4B,D).

### Pseudomonas can enhance outbreaks in large landscapes

In larger 100 × 100 landscapes, a greater percent of plants inhabited by *Pseudomonas* always reduces peak densities of aphids and parasitoids, but the effect of *Pseudomonas* on the timing and duration of these peaks varies by parasitoid responsiveness to aphid densities (Figure 4A,B). When parasitoids respond weakly to aphid densities, the peaks of both types of aphids last longer with greater percent *Pseudomonas* because the maximal parasitoid densities are reduced and delayed (Figure 4A). When parasitoids respond strongly to aphids, percent *Pseudomonas* does not change the timing or duration of peaks (Figure 4B).

**Figure 4:**
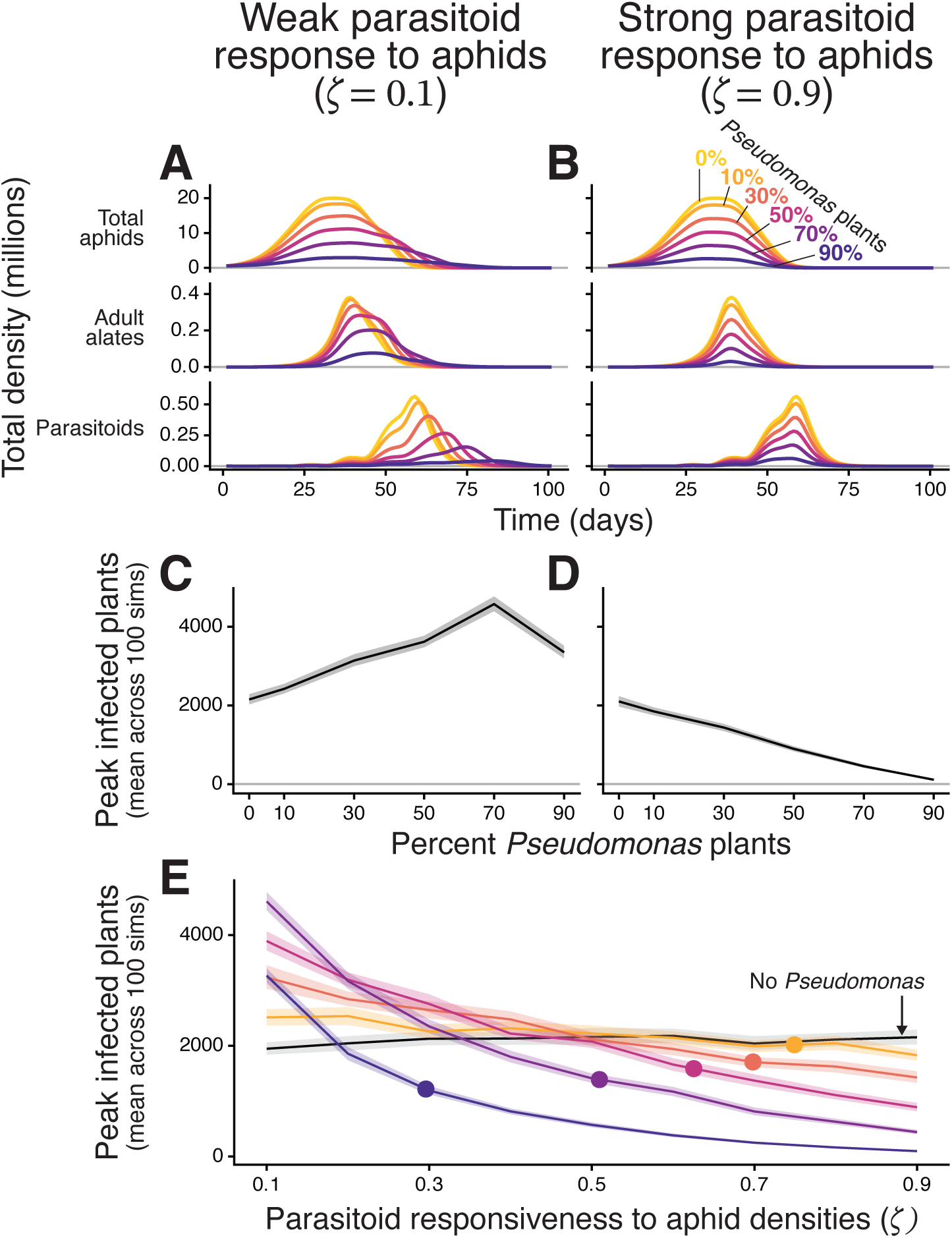
In larger 100 × 100 landscapes of pea plants, *Pseudomonas* occupancy affects aphid–parasitoid dynamics and virus outbreaks. Results are shown for (A,C) weak and (B,D) strong parasitoid responsiveness to aphid densities. (A,B) Time series of total aphids, adult alates, and adult parasitoids (females only) densities; color indicates *Pseudomonas*-inhabited plant percent. All densities are means through time across 100 simulations. (C,D) Peak infected plants versus percent *Pseudomonas* plants. (E) Peak infected plants versus parasitoid responsiveness. Color is the same as for A and B, except for *Pseudomonas*-free landscapes in black for easier comparison. Points indicate the estimate of *ζ* using empirical data from Ives et al. (1999). (C–E) Lines are means and ribbons are 95% confidence intervals across 100 simulations. Parameters and starting conditions are the same as for Figure 2, except that *Y*_0_ = 250.

The effects on virus transmission of increasing the percent of plants occupied by *Pseudomonas* coincides with results from our small landscapes: *Pseudomonas* promotes virus transmission when responsiveness is weak, and inhibits it when responsiveness is strong (Figure 4C,D). When parasitoid responsiveness is weak, however, the magnitude of the effect peaks at 70% *Pseudomonas* occupancy (Figure 4C). This non-monotonic relationship is because at very high (90%) *Pseudomonas* densities, the landscape-wide suppression of aphid abundances outweighs the effect of slower parasitoid population growth, as can be seen in the comparatively drastic reduction in adult alates that occurs between 70% and 90% *Pseudomonas* (Figure 4A). When parasitoid responsiveness is strong, virus transmission decreases consistently with greater *Pseudomonas* occupancy (Figure 4D).

### Empirical estimate of parasitoid responsiveness (ζ)

To estimate parasitoid responsiveness to aphid densities (*ζ*), we compared parasitoid densities on individual plants in our simulations to those predicted based on experimental tests of *A. ervi* parasitoid foraging (Ives et al. 1999). Because the empirically predicted parasitoid densities depend on the proportion of plants inhabited by *Pseudomonas*, we did so across a range of *Pseudomonas* occupancies. Our simulations match empirical predictions across a range of *ζ*, from 0.296 at 90% *Pseudomonas* occupancy to 0.749 at 10% *Pseudomonas* occupancy (Appendix S1: Figure S5). All empirical estimates of *ζ* fall within the range where *Pseudomonas* inhibits virus transmission, and the magnitude of this inhibiting effect strengthens with increasing *Pseudomonas* occupancy (Figure 4E).

## Discussion

Here we use a multi-scale model to show how *Pseudomonas* bacteria that protect individual plants from aphid vectors can promote aphid-borne virus transmission. We find that the host population-level impact of *Pseudomonas* depends on ecological interactions between aphids and their parasitoid natural enemies. Consequently, the effects of *Pseudomonas* protection on population-level virus transmission in plants can be fundamentally altered by a single key property of aphid–parasitoid interactions: the ability of parasitoids to track aphid densities. This study demonstrates that the effects of host protection can depend on the broader community context of vector-natural enemy interactions, addressing a gap in understanding how partial protection and trophic interactions jointly shape vector-borne disease dynamics (Okamoto and Amarasekare 2012, Wyckhuys et al. 2025). Specifically, we predict that when partial host protection generates spatial variation in vector populations that reduces natural enemy efficiency, it can undermine biocontrol and increase disease transmission. That effect is magnified when vectors increase dispersal in response to crowding.

A key takeaway from our study is that partial protection from vector-borne disease may not translate to population-level protection, due in large part to the strong ecological interactions between pea aphids and the parasitoid *A. ervi*. Such strong interactions between hosts and specialized parasitoids can generate unstable population cycles (Hassell 1978, Nicholson and Bailey 1935) that can undermine biocontrol. Many mechanisms can stabilize these dynamics by reducing the efficiency of parasitoid attacks, such as spatial clustering and invulnerable host stages (Hassell and May 1973, Murdoch et al. 2005, Murdoch and Oaten 1975). We find that *Pseudomonas* increases the spatial variability of aphid populations, which can reduce the efficiency of parasitoid attacks, slow parasitoid population growth, and decrease the amplitude of host–parasitoid cycles (Figure 4A). Examining longer-term dynamics of our multi-scale model would reveal whether partial protection of host populations could also stabilize otherwise ineffective biocontrol agents with high attack rates.

Consistent with past theory, parasitoid responsiveness emerges as a key parameter in our multiscale model. When we compared varying levels of parasitoid responsiveness in our model to results from an empirical study of parasitoid foraging times (Ives et al. 1999), the closest matches occurred for *ζ* values between 0.296–0.749 depending on *Pseudomonas* occupancy, which should typically result in *Pseudomonas* inhibiting virus transmission (Figure 4E). Since *A. ervi* is an aphid specialist, it should have a stronger response to aphid densities than most other natural enemies that have weaker and less tightly coupled ecological interactions with aphids and other vectors. A weaker response to aphid densities means that *Pseudomonas* should be more likely to promote virus transmission. However, generalist predators show less pronounced population responses to individual prey densities than specialists, so *Pseudomonas*-induced protection likely has a weaker effect on such predators. Extending our model framework to incorporate natural enemies that attack multiple prey species and show weaker aggregation and population growth in response to vector populations would help clarify how partial population protection of plants affects epidemiological outcomes across systems with different types of vector biocontrol agents.

While parasitoid responsiveness influences whether *Pseudomonas* promotes or inhibits simulated viral outbreaks, the magnitude of that effect varies with density-dependent alate production (Appendix S1: Figure S3). This is because it causes an additional link between aphid density and virus transmission: Greater plant-level aphid density increases the production of winged alates, the only morph capable of vectoring virus in our model. This additional feedback is partly why our results differ from other theoretical work showing that predator effects on vector density should not be as important as effects on vector behavior (Crowder et al. 2019). Dispersal polymorphisms are common across a wide range of insect species, including a number of important plant-virus vectors like aphids and leafhoppers that have density-dependent production of winged forms (Braendle et al. 2006, Roff 1986, Xu and Zhang 2017, Denno 1994). Density-dependent dispersal is also common more broadly, occurring in many organisms that lack distinct dispersing morphotypes (reviewed in Matthysen 2012). Thus, a key feature of our framework—feedbacks between local vector density and vector dispersal—is likely important for understanding vector-borne disease dynamics across many systems.

Although not included in our model, short-distance walking dispersal could also impact virus transmission. Aphids typically walk between plants after being dislodged by wind or rain, or after dropping to escape natural enemies (Fereres et al. 2017). Walking incurs costs through reduced feeding opportunities and desiccation risk (Dill et al. 1990), so long-distance walking is rare (Ben-Ari et al. 2015). Theoretical work has shown that when both non-winged aphids and alates can vector viruses at different spatial scales, maximal virus incidence occurs at moderate alate production, at least in the absence of parasitoids (Donnelly et al. 2019). The inclusion of parasitoids in models may be especially important for elucidating how walking dispersal affects virus transmission since parasitoids further promote walking in aphids (Hodge et al. 2011). Both long- and short-distance dispersal could impact disease dynamics, and our results suggest that these processes interact with parasitoids in ways that that could fundamentally alter predicted outcomes in field settings.

Distinct from vector dispersal, our model suggests that the dynamics of *Pseudomonas* spread through the landscape is likely to play an important role in outbreaks. *Pseudomonas* disperses through the water cycle (Morris et al. 2008) and potentially via pea aphid honeydew after infection (Stavrinides et al. 2009). However, no empirical studies have described the temporal dynamics of *Pseudomonas* dispersal or the extent to which this changes *Pseudomonas* densities in field conditions. Immigration order and priority effects often have strong effects on the assemblage of epiphytic microbial communities (Yang et al. 2023), so dispersal rates for *Pseudomonas* likely have complex, nonlinear relationships with *Pseudomonas* prevalence. Better descriptions of the spatiotemporal dynamics of *Pseudomonas* prevalence will be critical to understanding how *Pseudomonas* affects aphid-borne virus transmission because it dictates the extent to which (i) aphids always have spatial refuges from *Pseudomonas* infection and (ii) greater aphid densities increase *Pseudomonas* prevalence and future infection risk. *Pseudomonas* is also a phenotypically diverse genus with high variation across strains in their effects on aphid mortality and dispersal (Hendry et al. 2016, 2018, Herr et al. 2024, Smee et al. 2017, 2021, Smee and Hendry 2022) their ability to colonize and persist on plant surfaces (Smee et al. 2017), and their phytopathogenicity (Grenier et al. 2006). Understanding how these traits impact *Pseudomonas*–aphid–plant interactions will provide better quantitative predictions of how diverse communities of *Pseudomonas* impact biocontrol of aphid vectors.

When natural enemies respond weakly to vector densities, more protection provided to individual hosts (through greater *Pseudomonas* occupancy or aphid mortality, *ψ*) increases population-level virus transmission due to slowed population growth of natural enemies. This counterintuitive outcome demonstrates the utility of including realistic interactions between vectors and natural enemies in the simulation of vector-borne disease dynamics, and parallels other work finding that adding interactions within food webs can flip the direction of other species interactions (Fahimipour et al. 2025). Thus, the effectiveness of partial host population protection in suppressing disease transmission depends critically on the behavior of natural enemies.

## Acknowledgments

This work was supported by a National Institutes of Health training grant (project number 5T32AI145821-05) to LAN, and by the Cornell University College of Agriculture & Life Sciences (MAG).

## Author Contributions

Lucas Nell built the model and wrote the first draft of the manuscript. All other authors provided input in the model design and in editing the manuscript.

## Conflict of Interest Statement

The authors declare no conflicts of interest.

## Appendix S1

### S1: Host–parasitoid model

Because adult parasitoids are far more mobile than aphids, the single-field version of the model used here tracks adult, female parasitoids (*Y*) as one field-wide pool. To estimate aphid survival from parasitoid attack, we split these adult parasitoids across all *q* plants, and we allow parasitoid densities to increase with aphid density (Equation 1, repeated in Equation S7 below). In contrast to parasitoids, there are separate aphid and non-adult parasitoid populations for each plant. Aphids are stage-structured by both age (*n* days) and by morphology (winged “alates” and non-winged), so aphid populations are tracked using a length-2*n* vector of abundances by stage (**X**). Aphid stages in **X** are first ordered by morphology, then by age, so the first *n* stages are non-winged aphids, and the last *n* are alates. We also track stage-structured densities of parasitized, alive aphids (**P**, of length 7) and parasitized, dead aphids, known as mummies (**M**, of length 4).

For the aphid population in plant *i*, population growth follows the equation

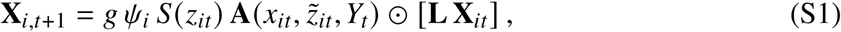

where *g* is survival from generalist predators and other miscellaneous sources of mortality, *ψ* is aphid survival from *Pseudomonas* infection (*ψ* = 1 on plants without *Pseudomonas*), *S*(·) is unparasitized aphid density dependence, *z* is total alive aphids 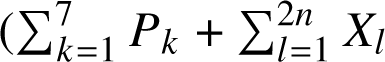, where *k* and *l* index by aphid stage), **A**(·) is a length-2*n* vector of the proportion of aphids surviving parasitism, *x* is total unparasitized aphids 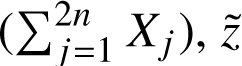, *z*° is relative alive aphid abundance 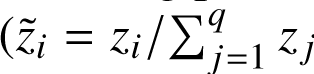, where *i* and *j* index through space), and ⊙ indicates element-wise multiplication (Hadamard product). **L** is the 2*n* × 2*n* matrix defining transitions among aphid stages. The exact structure of this matrix is described elsewhere (Equations S8, S10–S12; Nell et al. 2024), but the key elements are that aphid survival and reproduction vary by age, alates produce only non-winged offspring, and non-winged mothers can produce alate or non-winged offspring. Alates only producing non-winged offspring is based on previous empirical work (Nell et al. 2024, Christiansen-Weniger and Hardie 2000), as is the relationship between the density of aphids in a plant (*z*) and the proportion of offspring that are alates (*W* (·)):

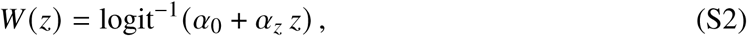

where *α*_0_ is the base proportion of alate offspring produced at aphid densities of zero, and *α_z_* is the effect of aphid density on alate production. We assume that parasitoids do not increase the production of alates in pea aphids, due to recent field observations suggesting that parasitism is unimportant for explaining alate production (Bosch and Ives 2023).

Parasitized aphids by stage in plant *i* (**P***_i_*) change according to

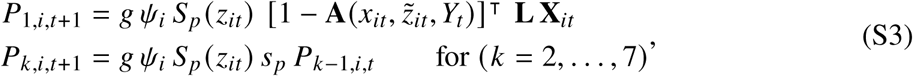

where *s_p_* is parasitized aphid daily survival *S_p_* (·) is parasitized aphid density dependence, and ⊺ indicates matrix transposition. The effects of density dependence for unparasitized and parasitized aphids increase with total aphids in the plant:

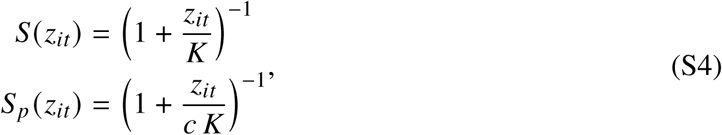

where *c* affects the density dependence for parasitized aphids relative to unparasitized (0 < *c* ≤ 1). Carrying capacity for unparasitized aphids on a given plant is *K* (*λgψ* − 1), where *λ* is the leading eigenvalue of **L**. Thus, *K* (*λ* − 1) represents the carrying capacity in the absence of *Pseudomonas* and generalist predators, and we manipulate carrying capacity by changing *K*.

Mummy (**M**) populations on plant *i* follow

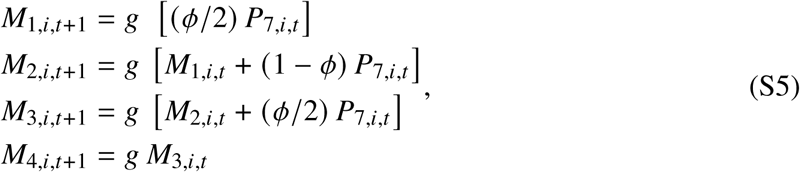

where *φ* is a number from 0 to 1 that smooths parasitoid development and *P*_7,*i*,*l*_ is the number of parasitized, living aphids on plant *i* in the stage immediately before transitioning to mummies.

Mummies across all *q* plants transition to the landscape-wide adult, female parasitoid population:

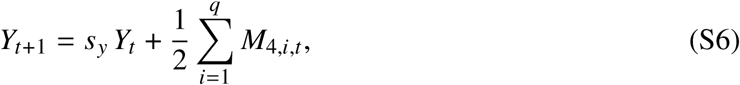

where *s_y_* is daily survival for adult parasitoids, *M*_4,*i*_ is density of last-stage mummies on plant *i*, and half of offspring are assumed to be female.

The density of adult, female parasitoids searching for aphid hosts on plant *i* (*γ*(·)) is affected by the relative number of aphids on that plant (*z*°*_i_*):

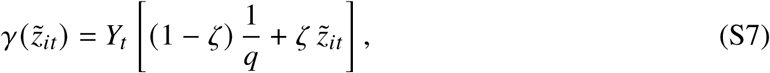

where *ζ* is a constant between 0 and 1 that determines the extent to which parasitoid densities are a linear function of aphid densities (Godfray and Pacala 1992). When *ζ* = 0, parasitoid densities are evenly distributed regardless of aphid densities, whereas when *ζ* = 1, there is a linear relationship between parasitoid and relative aphid densities.

Because parasitoid abundances vary across plants, so do the proportion of aphids that escape parasitoid attacks (**A**):

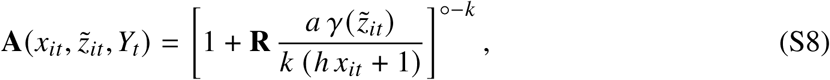

where **R** is a length-2*n* vector giving the relative parasitoid attack rates by aphid stage, *a* is the overall parasitoid attack rate, ℎ is the parasitoid attack-rate handling time, *k* is the negative binomial distribution’s aggregation parameter, and ^◦^ indicates element-wise power (Hadamard power).

### S2: Large plot dynamics are mildly sensitive to *Pseudomonas* repellence and viral attraction

When we allow virus infection and *Pseudomonas* presence to affect plant attractiveness to alates, the probability that a dispersing alate will choose plant *j* when they are on plant *i* is

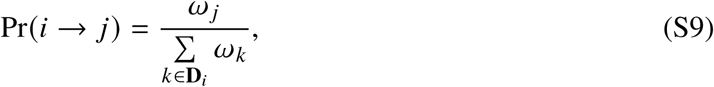

where *ω_j_* is the sampling weight of plant *j*, and **D***_i_* is a vector of indices for all plants within *r*_fly_ of plant *i* that does not include *i*. Sampling weights depend only on the state of the plant:

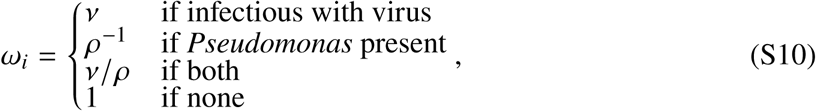

where *ν* is the attractant effect of the virus, and *ρ* is the repellent effect of *Pseudomonas*; *ν* ≥ 1 and *ρ* ≥ 1.

The effect of *Pseudomonas* repellence of alates (*ρ* = 5) depends on whether the virus starts on a *Pseudomonas*-inhabited plant. When the virus starts on an uninhabited plant, *Pseudomonas* repellence increases virus transmission (Figure S6E,G). This effect increases in strength with *Pseudomonas* prevalence and occurs because *Pseudomonas* repellence causes the initial infectious plant, being uninhabited by *Pseudomonas*, to receive greater numbers of incoming alates because it is surrounded by plants that repel them. *Pseudomonas* repellence promoting virus transmission makes the relationship between peak infected plants and *Pseudomonas* occupancy more concave and causes *Pseudomonas* occupancy to have a greater total positive effect on peak infected plants. In contrast, when the virus starts on a *Pseudomonas*-inhabited plant, *Pseudomonas* repellence decreases virus transmission (Figure S6F,H) because of the repellent effect on alates visiting the initially infectious plant. The inhibition of virus transmission caused by *Pseudomonas* repellence occurs most strongly when *Pseudomonas* is present on the landscape but at low density because the initially infectious plant is especially repellent to alates in comparison to its mostly non-repellent neighbors. *Pseudomonas* can have a net inhibitory effect on virus transmission at low *Pseudomonas* occupancy (compared to *Pseudomonas*-free landscapes) even when parasitoid responsiveness is weak, provided the virus starts on a *Pseudomonas*-inhabited plant (Figure S6F).

Viruses attracting alates (*ν* = 5) consistently results in greater virus transmission (Figure S6I–L). When parasitoids respond weakly to aphid densities, the effect of virus attraction is relatively constant across *Pseudomonas* occupancy (Figure S6I,J). When parasitoids respond strongly to aphid densities, virus attraction increases virus transmission less as *Pseudomonas* occupancy increases (Figure S6K,L) because the effect of *Pseudomonas* in inhibiting virus transmission overwhelms any effect of virus attraction. Consequently, virus attraction causes increasing *Pseudomonas* occupancy to have a stronger absolute effect in inhibiting virus transmission (Figure S6K,L). Virus attraction has the greatest effect on virus transmission when the virus starts on a *Pseudomonas* inhabited plant (Figure S6J,L) because those plants produce few alates, so incoming alates are necessary for the virus to pass from them to other plants.

Virus attraction consistently promotes virus transmission, even when virus outbreaks are large, due to three features of our model: explicit spatial structure, limited vector dispersal, and multiple visits for each alate vector flight. Spatial structure of host plants paired with limited alate dispersal distance results in a traveling wave of infection spreading across the landscape. Because virus prevalence is low at the front of this wave, virus attraction has a net positive effect on virus transmission here. Alates visiting multiple plants per dispersal (E(new plants visited) = 5 for *w* = 0.2 used throughout) amplifies virus transmission at the wave front because even when they are drawn to an infected plant, they typically leave to probe nearby uninfected ones. These elements together cause virus attraction to more effectively push the wave of infection across the landscape (Figure S7). Although virus attraction reduces virus transmission in areas behind the traveling wave, a faster wave takes better advantage of the limited time where alates are abundant, typically resulting in more infected plants.

In other general models of vector-borne disease (McElhany et al. 1995, Kingsolver 1987, Sisterson 2008) and models of aphid-vectored viruses (Donnelly et al. 2019), vector attraction to infectious hosts causes infectious hosts to act as sinks for vector visits when many hosts are infected, which suppresses large disease outbreaks. For our model, alate vectors being attracted to virus-infected plants consistently promotes virus transmission, even when virus prevalence is high. This occurs because in our spatially explicit model where alates can only disperse to nearby plants, virus infection moves through the landscape as a traveling wave. Attraction of alates to infected plants accelerates this wave of infection because (i) virus prevalence is low at the front of the wave and (ii) alates visit multiple host plants each time they disperse (Figure S7). Our results mirror those from other spatial models showing that vector attraction to infected hosts can increase the speed of traveling waves (Chamchod and Britton 2011, Hamelin et al. 2023). In our system where plants stay infected with the virus indefinitely, a faster wave of infection usually means more plants infected throughout the growing season. A number of empirical examples show that viruses can manipulate aphid vectors into preferentially probing, then leaving infected plants (Carmo-Sousa et al. 2016, Mauck et al. 2010), and this should increase virus transmission (Donnelly et al. 2019). We expect that having virus-infected plants attract, then repel, alates would increase virus transmission in our model and have strong interactions with *Pseudomonas* repellence and the type of initially virus-infected plant. Our results suggest that however complicated, virus attraction will not change the key result that *Pseudomonas*-induced protection can exacerbate population-level virus outbreaks.

### S3: Other factors have negligible effects in large landscapes

We manipulated a number of factors in our large-landscape simulations that did not appreciably change our results. First, we reduced loading probabilities (*δ_a_* = *δ_p_* = 0.05 instead of 0.5) to cause outbreaks (at least one new infection) to not always occur. The results mirror those from our default simulations with greater loading probabilities (shown in Figure 4A,B), the exceptions only occurring when disease outcomes approached their minimum or maximum possible values (Figure S8). We also manipulated host-plant attractiveness and variability in starting aphid densities (as in Figure S6) with reduce loading probabilities, in which case we also found similar results; the only exception was that including variability in aphid densities (*σ_N_* = 50) had a smaller effect (Figure S9).

We also spatially clustered *Pseudomonas* locations. To produce clustered patterns of *Pseudomonas* locations, we used iterative weighted sampling. To start, all locations were weighted equally, then each *Pseudomonas* location sampled caused the sample weights for all neighboring locations to be multiplied by *τ*; we used *τ* = 3 for all clustered landscapes. Spatially clustering *Pseudomonas* locations caused virtually no effect on virus transmission (Figure S10).

Lastly, we located the plant initially infectious with the virus on the interior of the landscape (*x* = 50, *y* = 50, versus *x* = 1, *y* = 1). When the initially infectious plant started on the interior, more plants are infected overall because there are more alate-producing and uninfected plants surrounding that plant. However, neither the effect of *Pseudomonas* nor the effects of other manipulated factors changed substantially (Figure S11).

### S4: Small plot dynamics are sensitive to initial insect densities and aphid carrying capacity

Although starting densities of aphids (*N*_0_) and parasitoids (*Y*_0_) can reduce or eliminate the effect of *Pseudomonas*, they never cause *Pseudomonas* to reverse its effect on virus transmission based on parasitoid responsiveness. Across the initial densities we simulated, *Pseudomonas* presence can increase the mean number of peak infected plants by up to 5.18 (Figure S12E) and decrease it by up to 2.70 (Figure S12F) for weak and strong parasitoid responsiveness, respectively. Regardless of parasitoid responsiveness to aphid densities, effects of changing the initial insect densities on virus transmission are often strong enough to overwhelm effects of *Pseudomonas* presence (Figure S12). If initial parasitoid densities are too high relative to aphid densities (*Y*_0_/*N*_0_ ≤ ∼0.03), parasitoids suppress alate production resulting in no new virus infections (Figure S13A,B). If initial parasitoid densities are too low (*Y*_0_/*N*_0_ ≥ ∼0.01), aphids are released from parasitism for long enough that all plants get infected before parasitoids recover (Figure S13C,D).

Increasing aphid carrying capacity (*K* (*λ* − 1)) increases virus transmission since it allows for greater aphid densities, but it does so non-uniformly across parasitoid responsiveness and *Pseudomonas* presence. When parasitoids respond weakly to aphids (*ζ* = 0.1), increasing the aphid carrying capacity causes a greater increase in virus transmission for *Pseudomonas*-inhabited landscapes than for landscapes without *Pseudomonas* (Figure S14A,C,E). Doing so when parasitoids respond strongly to aphids (*ζ* = 0.9) causes virus transmission to be promoted more strongly for non-*Pseudomonas* landscapes These changes arise because increasing aphid carrying capacity allows plants that already have the largest aphid populations to support even more aphids. The end result is that increasing aphid carrying capacity amplifies whatever effect *Pseudomonas* already has based on parasitoid responses to aphids.

### S5: Variability in initial aphid densities exaggerates the effects of *Pseudomonas* on virus transmission

For larger landscapes, we conducted a series of simulations that included variation among plants in their starting aphid densities by generating starting densities from a lognormal distribution. In this case, *N*_0_ and *σ_N_* are the mean and standard deviation of the resulting distribution, not of the underlying normal distribution. The parameters used for the underlying normal distribution were

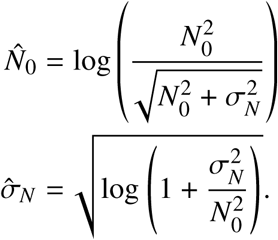

When initial aphid densities vary across plants (*σ_N_* = 50), it magnifies the effects of *Pseudomonas* on virus transmission. When parasitoid responsiveness is weak (*ζ* = 0.1), variable initial aphid densities increase peak infected plants as *Pseudomonas* occupancy increases (Figure S6A,B). This happens because variable initial aphid densities adds additional spatial variability to aphid densities that further slows parasitoid population growth. When parasitoid responsiveness is strong (*ζ* = 0.9), variable initial aphid densities reduce peak infected plants (Figure S6C,D) because nonlinearity causes the mean trajectory to be lower than when starting from a fixed density (Figure S15). This effect is less apparent at high *Pseudomonas* occupancy because all *Pseudomonas*-inhabited plants have similar (near zero) aphid densities through time.

### S6: Virus transmission is suppressed when the virus starts on a *Pseudomonas*-inhabited plant

When the virus starts in a plant protected by *Pseudomonas*, *Pseudomonas*-induced aphid mortality reduces the number of alates leaving, hindering virus transmission (black lines in Figure S6A–D). When parasitoids respond weakly to aphid densities, this effect is strongest when *Pseudomonas* is very common on the landscape (90% occupancy; black lines in Figure S6A versus B). This causes the net effect of very high *Pseudomonas* occupancy (when compared to no *Pseudomonas*) to switch from promoting to inhibiting virus transmission despite weak parasitoid responsiveness, although *Pseudomonas* still has a virus-promoting effect below 90% occupancy. The stronger effect of virus starting location at higher *Pseudomonas* occupancy happens because as *Pseudomonas* becomes more common, there are fewer alates being generated on the landscape that can immigrate into the initially infectious plant to transmit the virus to others. Viruses on *Pseudomonas*-inhabited plants are especially reliant on incoming alates to transmit to other plants because nearly zero alates are being generated on those plants. When parasitoids respond strongly to aphid densities, the virus starting on a *Pseudomonas*-inhabited plant has a consistent negative effect on peak infected plants, except at higher *Pseudomonas* occupancies when numbers of peak infected plants approach the minimum possible value of 2 (black lines in Figure S6C versus D).

### S7: Parameterizing model and software

We created our simulation software using a combination of R version 4.5.2 (R Core Team 2025) and C++ using the Rcpp (Eddelbuettel 2013) package.

Parameter values for the host–parasitoid model were based on those from previous work (Meisner et al. 2014, Nell et al. 2024) (Table S1B). We chose values for our aphid transition matrix (**L**) based on the aphid clone with no resistance to parasitism from Nell et al. (2024).

We used a mixed approach to our spatial and disease model parameters. For small landscapes, we used an alate dispersal radius (*r*_fly_) of 1 to avoid all alates being able to disperse to all other plants. For more realistic, larger landscapes, we parameterized our fly radius based on field data from Nemecek (1993), who found that aphid dispersal distances (in meters) followed a Weibull distribution with shape 0.6569 and scale 9.613. We used 7.34 for our radius parameter, which is the median of this Weibull distribution divided by 0.75. We divided by 0.75 to convert from meters to plants since 0.75 meters is a typical row spacing in fresh pea crops (Phillips et al. 2025). For the probability that an alate stays on a plant indefinitely (*w*), we used 0.2 as by Donnelly et al. (2019). When simulating small landscapes, we used 0.5 for the probabilities of viruses loading onto plants and aphids (*δ_a_*, *δ_p_*) because this resulted in non-extreme numbers of infected plants. Because larger landscapes had more total virus-vectoring alates, simulations with these loading probabilities always had outbreaks (at least one new plant being infected). Therefore, we also conducted large-landscape simulations where *δ_a_* = *δ_p_* = 0.05 to show disease outcomes when not all simulations had outbreaks. For parameters related to plant attractiveness (*ν* and *ρ*) we compared outcomes when these factors had no effect (*ν* = 1 and *ρ* = 1) to when they had a strong effect (*ν* = 5 or *ρ* = 5). For the remaining parameters (*ψ* and *ζ*), we chose values that resulted in *Pseudomonas* inhibiting or promoting disease spread to about the same degree. We then varied each parameter to test for how sensitive our outcomes were to their values.

**Table S1:**
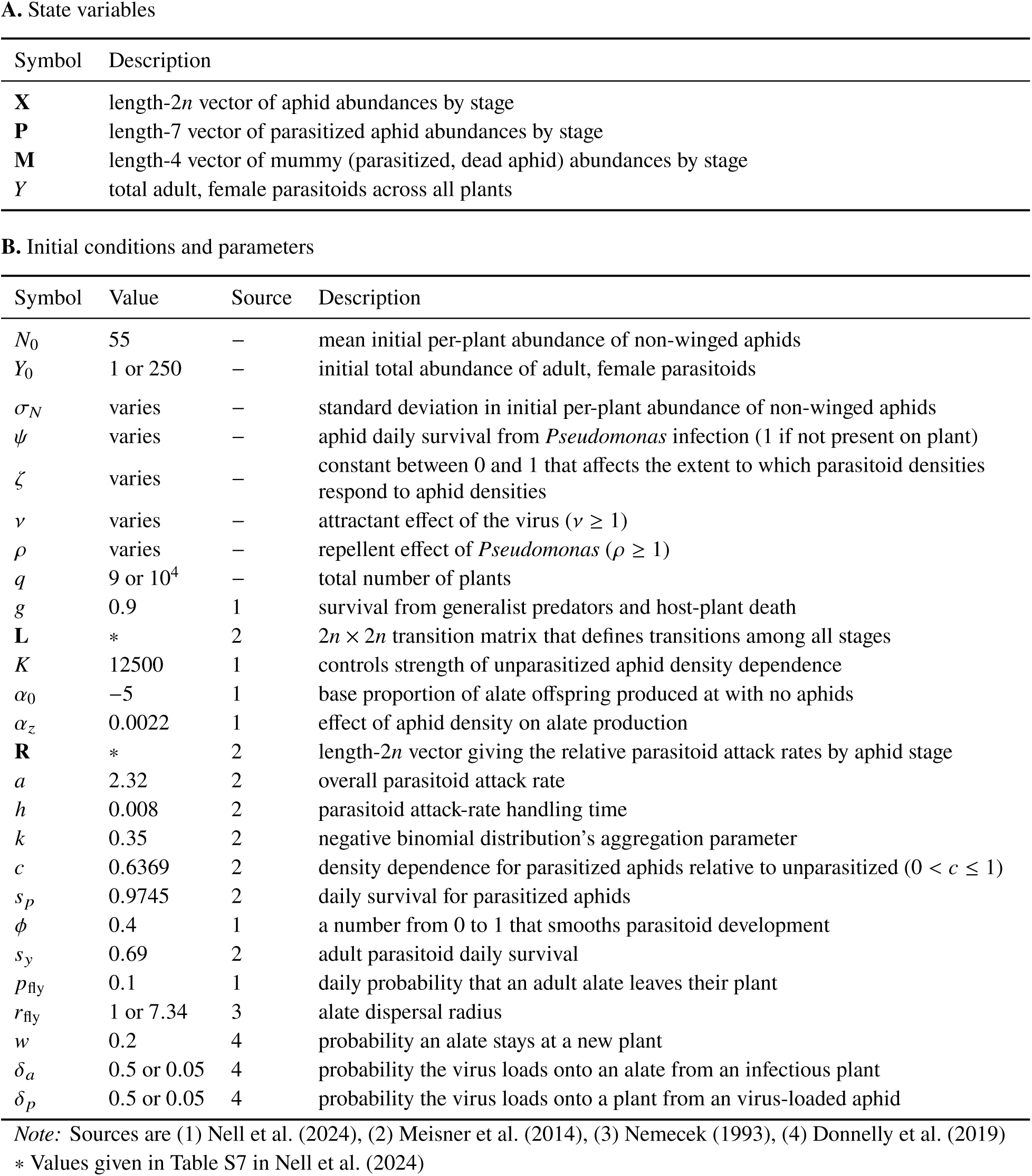
Model (A) state variables, as well as (B) initial conditions and parameter values with sources. Multiple values indicate those for small- and large-landscape simulations, respectively.

| Symbol | Description |
| --- | --- |
| **X** | length-$2n$ vector of aphid abundances by stage |
| **P** | length-7 vector of parasitized aphid abundances by stage |
| **M** | length-4 vector of mummy (parasitized, dead aphid) abundances by stage |
| **Y** | total adult, female parasitoids across all plants |

**B. Initial conditions and parameters**
| Symbol | Value | Source | Description |
| --- | --- | --- | --- |
| $N_0$ | 55 | – | mean initial per-plant abundance of non-winged aphids |
| $Y_0$ | 1 or 250 | – | initial total abundance of adult, female parasitoids |
| $\sigma_N$ | varies | – | standard deviation in initial per-plant abundance of non-winged aphids |
| $\psi$ | varies | – | aphid daily survival from *Pseudomonas* infection (1 if not present on plant) |
| $\zeta$ | varies | – | constant between 0 and 1 that affects the extent to which parasitoid densities respond to aphid densities |
| $\nu$ | varies | – | attractant effect of the virus ($\nu \geq 1$) |
| $\rho$ | varies | – | repellent effect of *Pseudomonas* ($\rho \geq 1$) |
| $q$ | 9 or $10^4$ | – | total number of plants |
| $g$ | 0.9 | 1 | survival from generalist predators and host-plant death |
| **L** | * | 2 | $2n \times 2n$ transition matrix that defines transitions among all stages |
| $K$ | 12500 | 1 | controls strength of unparasitized aphid density dependence |
| $\alpha_0$ | –5 | 1 | base proportion of alate offspring produced at with no aphids |
| $\alpha_z$ | 0.0022 | 1 | effect of aphid density on alate production |
| **R** | * | 2 | length-$2n$ vector giving the relative parasitoid attack rates by aphid stage |
| $a$ | 2.32 | 2 | overall parasitoid attack rate |
| $h$ | 0.008 | 2 | parasitoid attack-rate handling time |
| $k$ | 0.35 | 2 | negative binomial distribution's aggregation parameter |
| $c$ | 0.6369 | 2 | density dependence for parasitized aphids relative to unparasitized ($0 < c \leq 1$) |
| $s_p$ | 0.9745 | 2 | daily survival for parasitized aphids |
| $\phi$ | 0.4 | 1 | a number from 0 to 1 that smooths parasitoid development |
| $s_y$ | 0.69 | 2 | adult parasitoid daily survival |
| $p_{fly}$ | 0.1 | 1 | daily probability that an adult alate leaves their plant |
| $r_{fly}$ | 1 or 7.34 | 3 | alate dispersal radius |
| $w$ | 0.2 | 4 | probability an alate stays at a new plant |
| $\delta_a$ | 0.5 or 0.05 | 4 | probability the virus loads onto an alate from an infectious plant |
| $\delta_p$ | 0.5 or 0.05 | 4 | probability the virus loads onto a plant from an virus-loaded aphid |

*Note:* Sources are (1) Nell et al. (2024), (2) Meisner et al. (2014), (3) Nemecek (1993), (4) Donnelly et al. (2019)
\* Values given in Table S7 in Nell et al. (2024)

### Supplemental Figures

**Figure S1:**
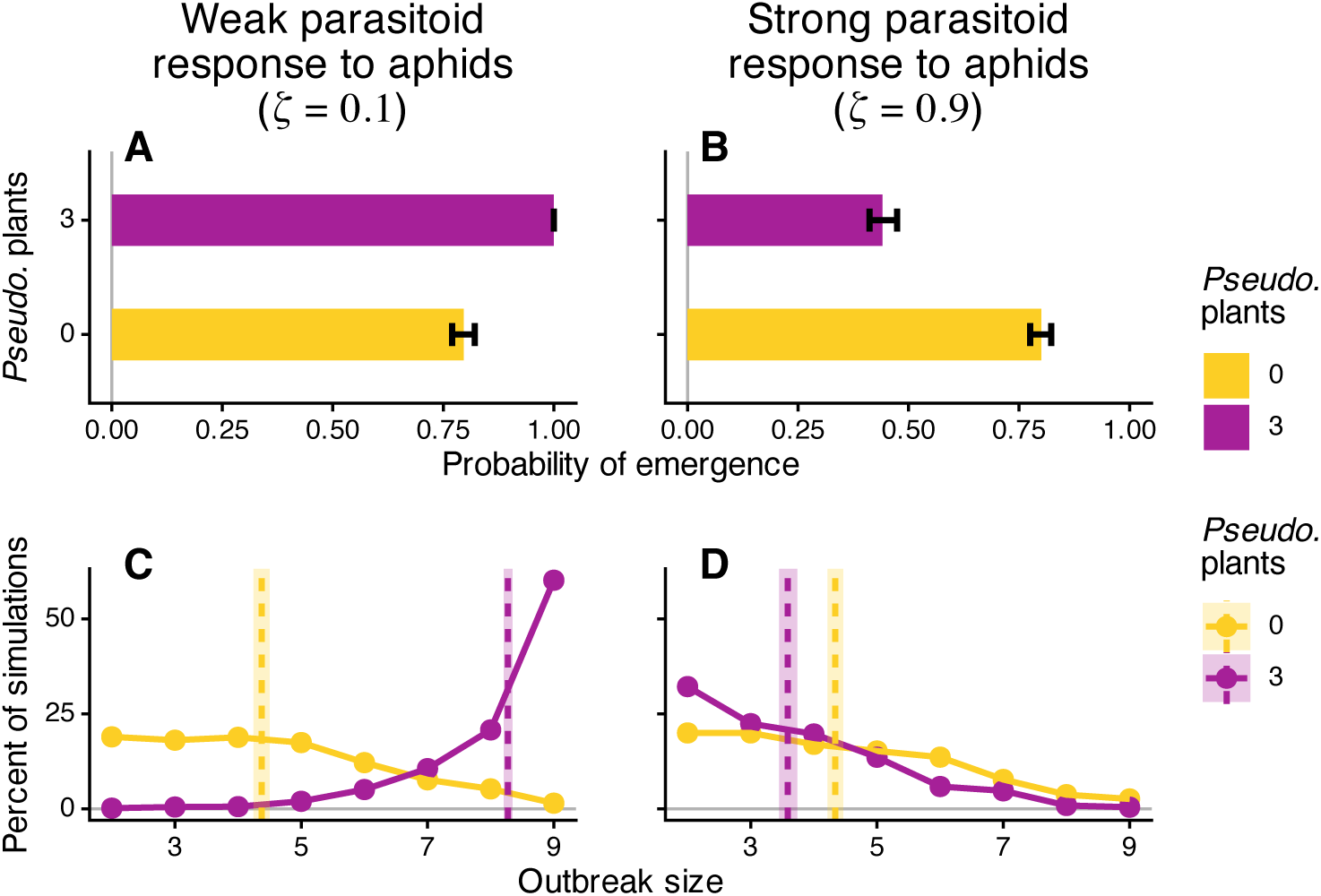
In 3 × 3 landscapes of pea plants, *Pseudomonas* can either promote virus outbreaks by slowing the population growth of parasitoid wasps or inhibit them by killing aphids. These disparate outcomes occur when parasitoids respond (A,C) weakly or (B,D) strongly to aphid densities, respectively. In all plots, color indicates whether landscapes contained (magenta) 3 or (gold) 0 *Pseudomonas*-inhabited plants. (A,B) Bar plots show the probability of emergence (defined as at least one new plant infected); error bars indicate 95% confidence intervals. (C,D) Frequency plots show the distribution of outbreak sizes (when an outbreak occurred); dashed vertical lines indicate means, and vertical ribbons indicate 95% confidence intervals. Simulations used the same parameters and starting conditions as in Figure 2.

**Figure S2:**
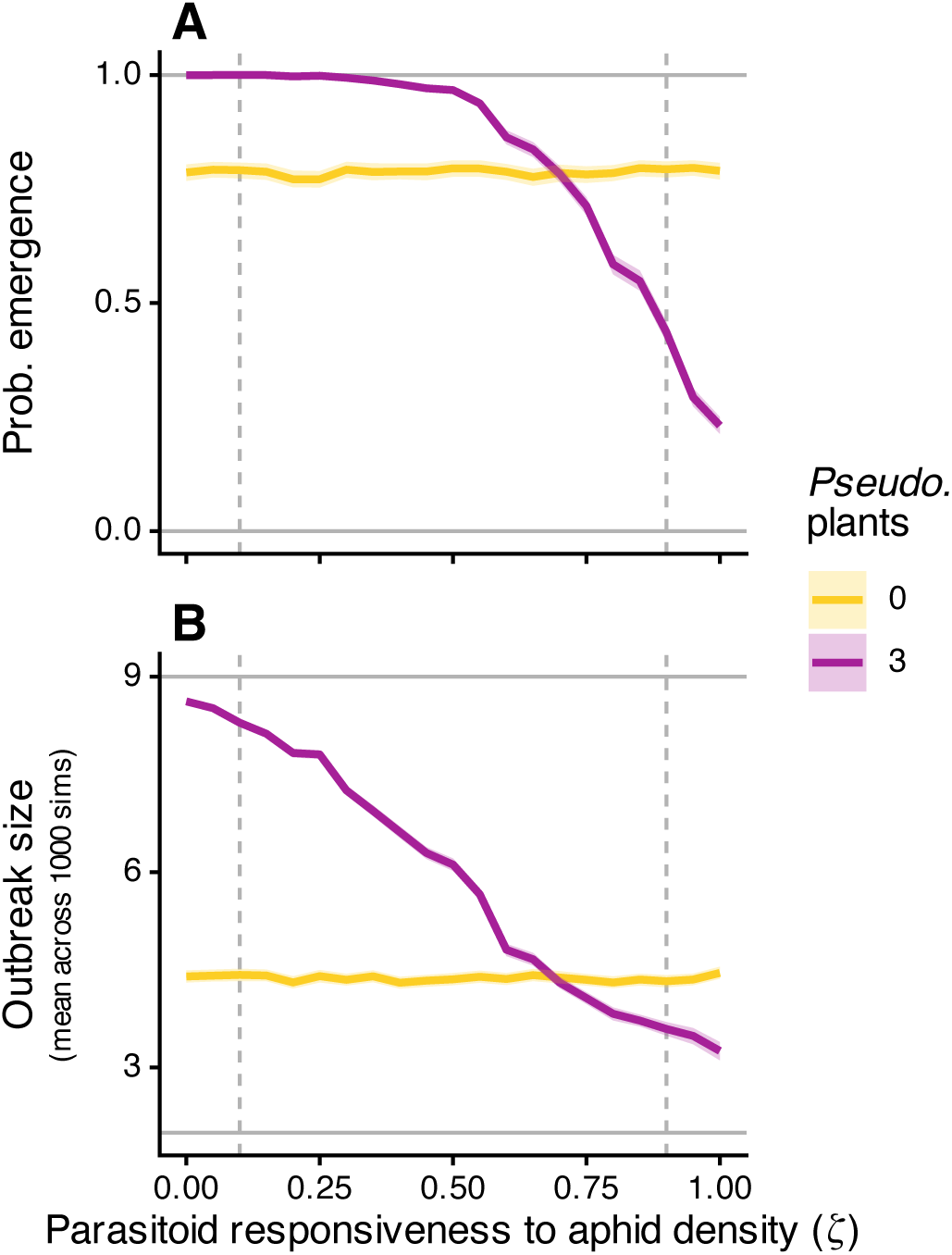
Line graphs showing the (A) probability of emergence and (B) mean outbreak size across values of parasitoid responsiveness to aphid densities (*ζ*). Lines are shown for landscapes (magenta) with and (gold) without *Pseudomonas*-inhabited plants. Vertical dashed lines indicate the *ζ* values used for the original simulations shown in Figure 2. Shaded ribbons indicate 95% confidence intervals.

**Figure S3:**
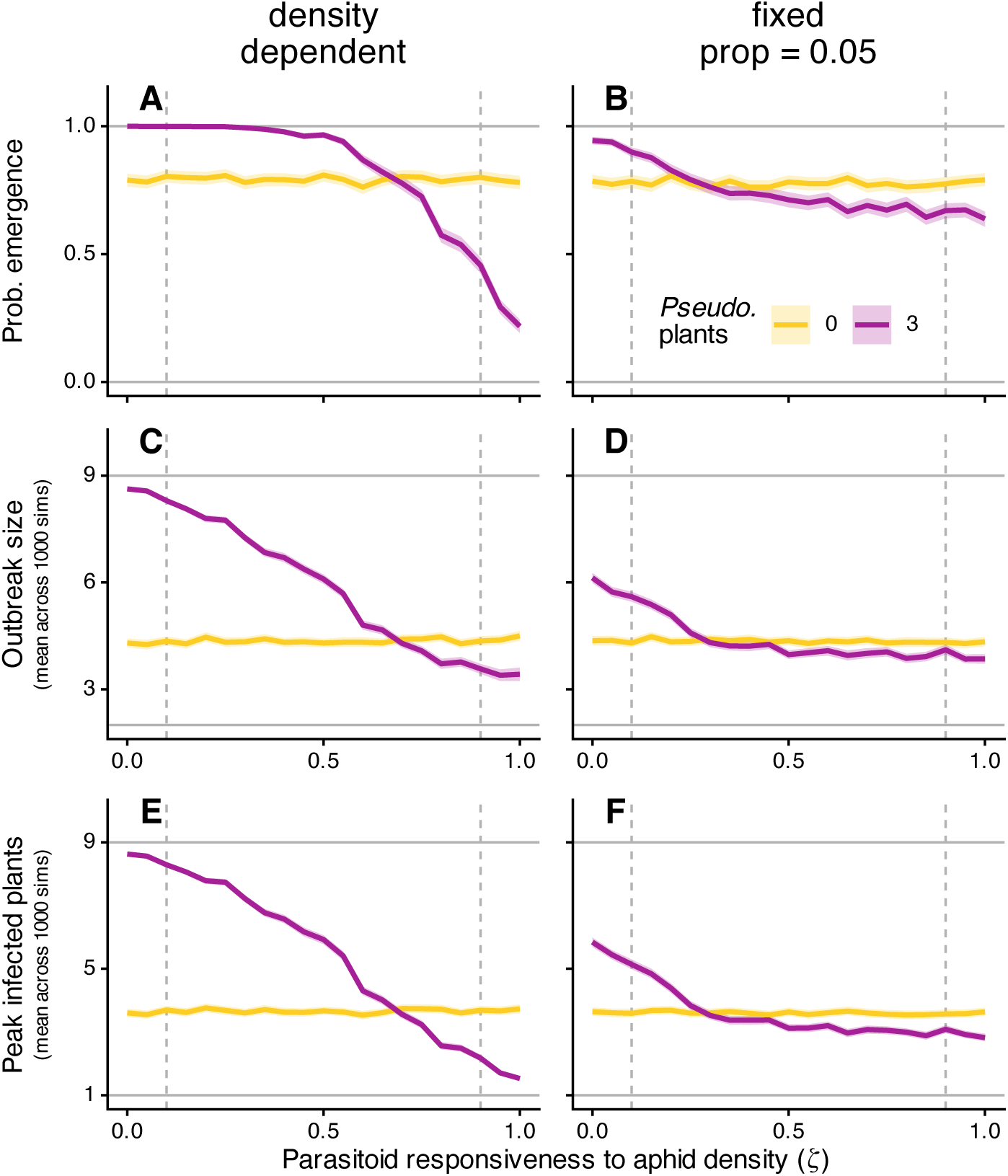
Effects of density-dependent dispersal on the relationship between virus outcomes and parasitoid responsiveness to aphid densities (*ζ*) in small 3 × 3 landscapes. Line graphs show the (A,B) probability of emergence, (C,D) mean outbreak size, and (E,F) mean peak infected plants across values of parasitoid responsiveness. Lines are shown for landscapes (magenta) with and (gold) without *Pseudomonas*-inhabited plants. Vertical dashed lines indicate the parasitoid responsiveness values used for the original simulations shown in Figure 2. Panel columns separate the type of alate production: (A,C,E) density-dependent alate proportion or (B,D,F) fixed alate proportion of 0.05. We chose this fixed value because it resulted in similar outbreak measures for *Pseudomonas*-free landscapes. Density-dependent alate proportion is the default used throughout and produces a proportion of offspring that are alates (*W*) based on Equation S2 with *α*_0_ = −5 and *α_z_* = 0.0022, which are based on results from Nell et al. (2024). For the fixed alate proportion, we set *α_z_* = 0 to cause alate production to no longer depend on aphid density, and we set *α*_0_ = logit(0.05) to achieve a fixed alate proportion of 0.05. All other parameters and starting conditions match Figure 2.

**Figure S4:**
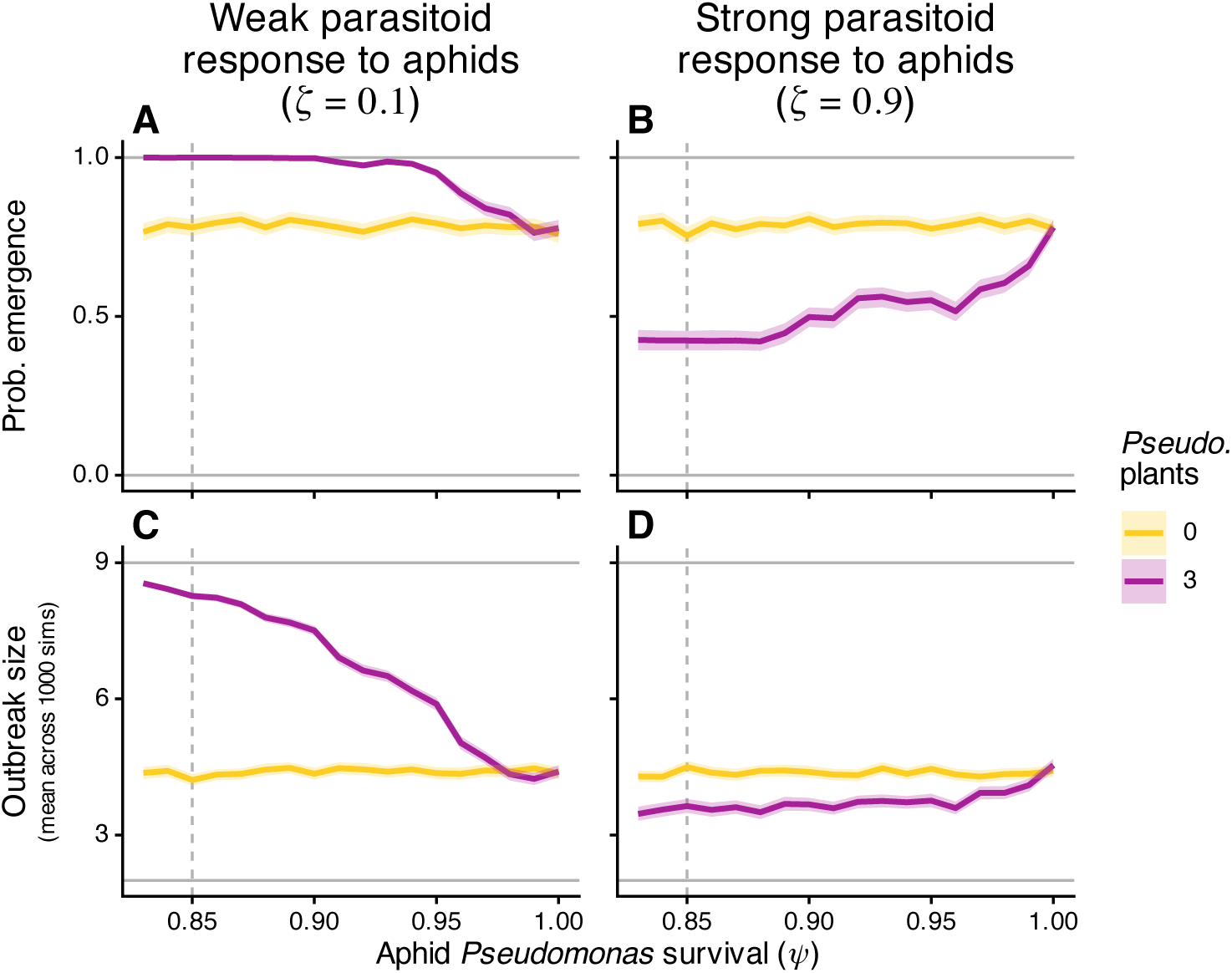
Line graphs showing the (A,B) probability of emergence and (C,D) mean outbreak size across values of aphid *Pseudomonas* survival (*ψ*), for parasitoids responding (A,C) weakly (*ζ* = 0.1) or (B,D) strongly (*ζ* = 0.9) to aphid densities. Lines are shown for landscapes (magenta) with and (gold) without *Pseudomonas*-inhabited plants. Vertical dashed lines indicate the *ψ* values used for the original simulations shown in Figure 2. Shaded ribbons indicate 95% confidence intervals.

**Figure S5:**
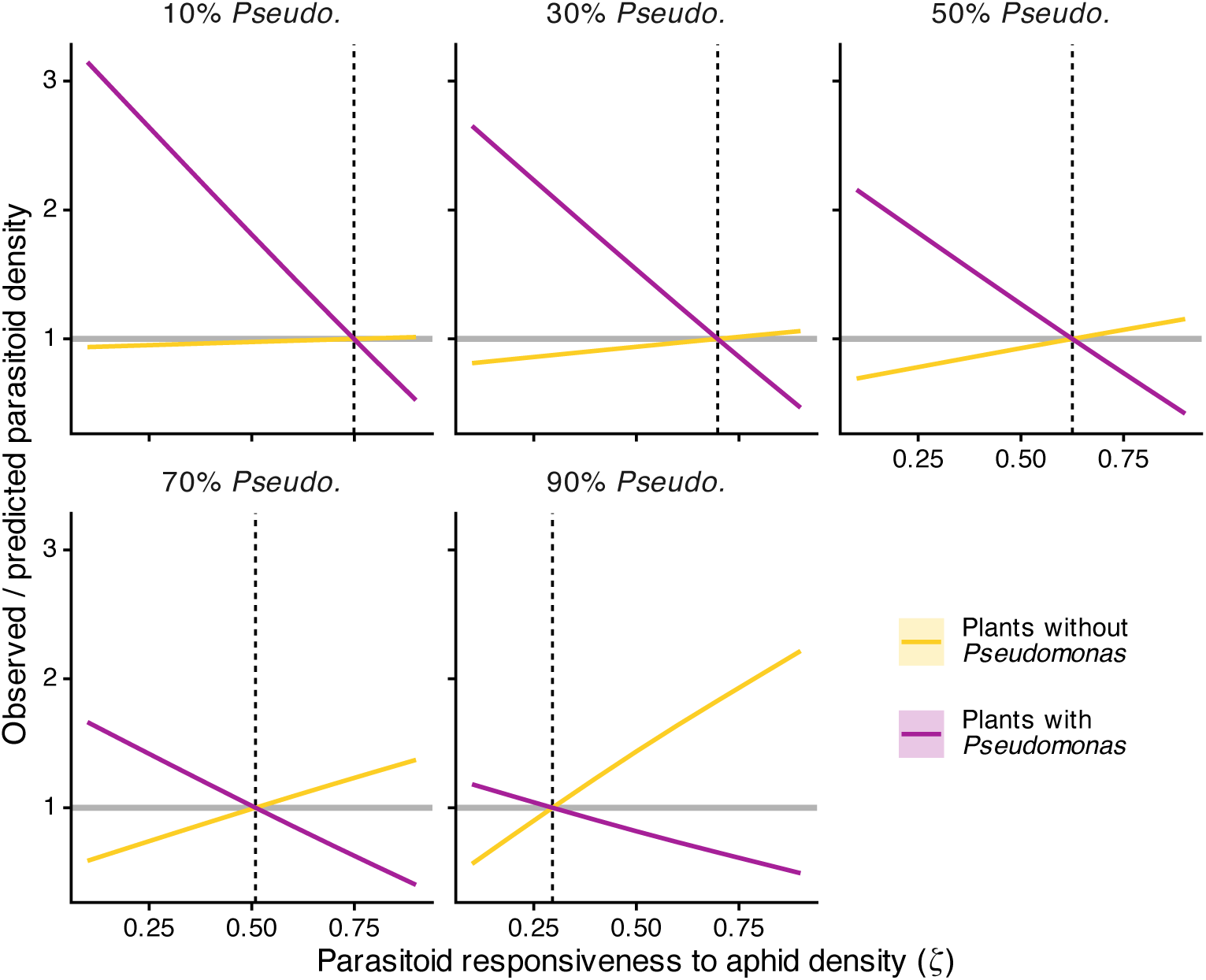
In Ives et al. (1999), they found that parasitoids spent about 3.76 more time foraging at plants where they encountered an aphid. For this figure, we simulated our model with large landscapes, and we varied (1) parasitoid responsiveness to aphid density (*ζ*) and (2) the number of plants inhabited by *Pseudomonas*. We then compared the observed parasitoid densities at the maximum total aphid density along the time series—when spatial variation in aphid densities is at its maximum—to parasitoid densities predicted based on Ives et al. (1999). For the predicted parasitoid densities, we assumed that parasitoids never encounter an aphid on a *Pseudomonas*-inhabited plant but always do on plants without *Pseudomonas*. Panels separate the percent of plants inhabited by *Pseudomonas*. Color indicates whether plants were inhabited by *Pseudomonas*. The y-axis is observed (in our simulations) divided by predicted (based on Ives et al. (1999)) parasitoid density, and the horizontal line indicates one. The x-axis is the value of *ζ*. Shaded ribbons indicate the 95% confidence intervals calculated using nonparametric bootstrapping across plants and the 12 repetitions per combination of *ζ* and % *Pseudomonas*. These ribbons are usually not visible because they are so narrow. The vertical dashed line is the value of *ζ* where observed / predicted parasitoid densities equal 1 (i.e., observed = predicted); we computed this using linear interpolation and one-dimensional root-finding (R functions approxfun and uniroot, respectively). Parameters other than *ζ* correspond to those used in the simulations in Figure 4.

**Figure S6:**
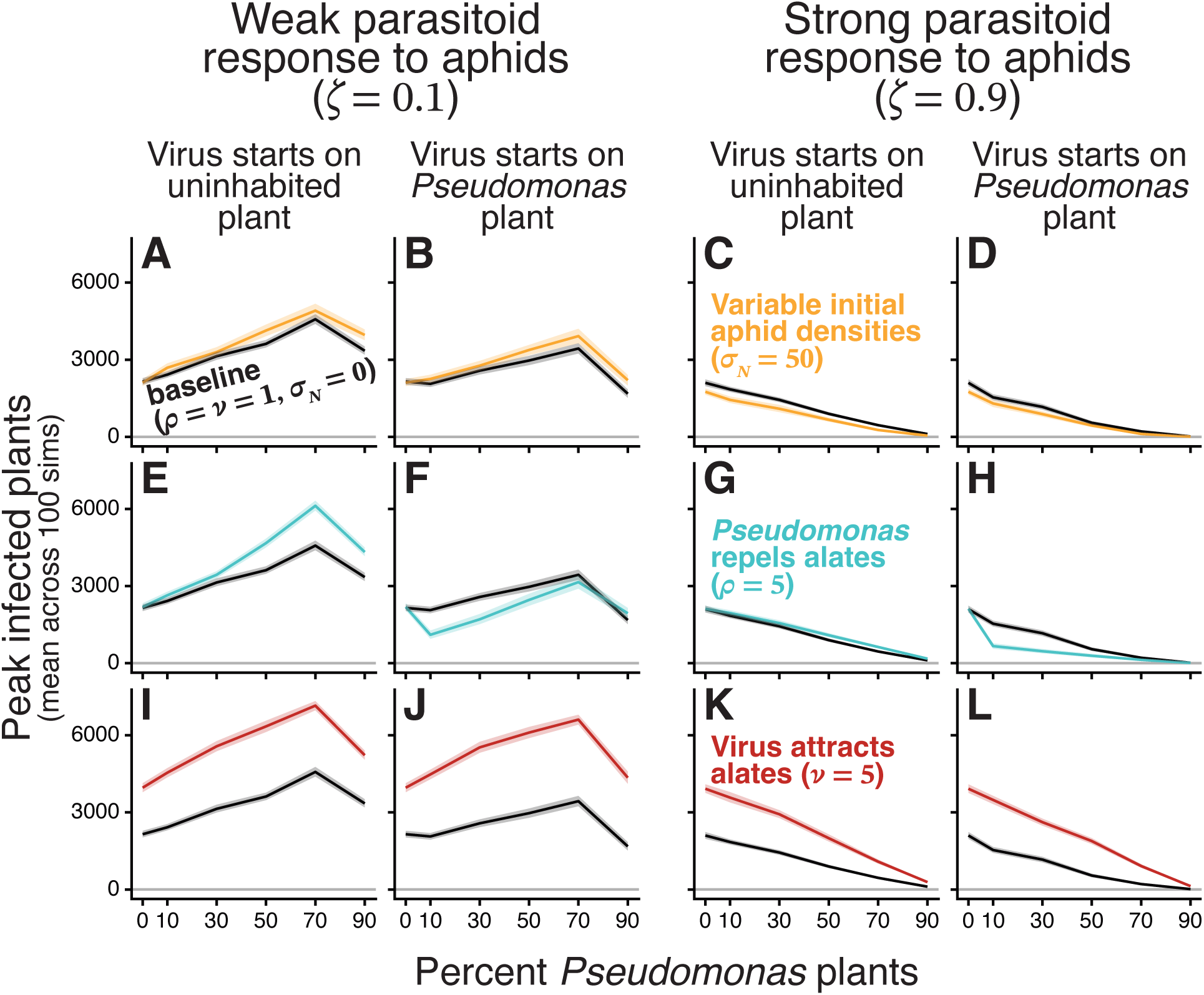
In larger 100 × 100 landscapes of pea plants, the effect of *Pseudomonas* on outbreaks can be altered by factors affecting the spatial distribution of insects and viruses. Results are shown for (A,B,E,F,I,J) weak and (C,D,G,H,K,L) strong parasitoid responsiveness to aphid densities, and for whether the virus starts on an (A,C,E,G,I,K) uninhabited or (B,D,F,H,J,L) *Pseudomonas*-inhabited plant. Line plots show mean peak infected plants across 100 simulations, versus percent *Pseudomonas* plants. Shaded ribbons show 95% confidence intervals. In all panels, the black line is results from the baseline simulations where attractiveness to alates and starting aphid densities are uniform across plants. Each colored line is from simulations where either (A–D) plants vary in starting aphid densities (*σ_N_* = 50), (E–H) *Pseudomonas* repels alates (*ρ* = 5), or (I–L) virus infection attracts alates (*ν* = 5). Other than those indicated, parameters are the same as for Figure 4.

**Figure S7:**
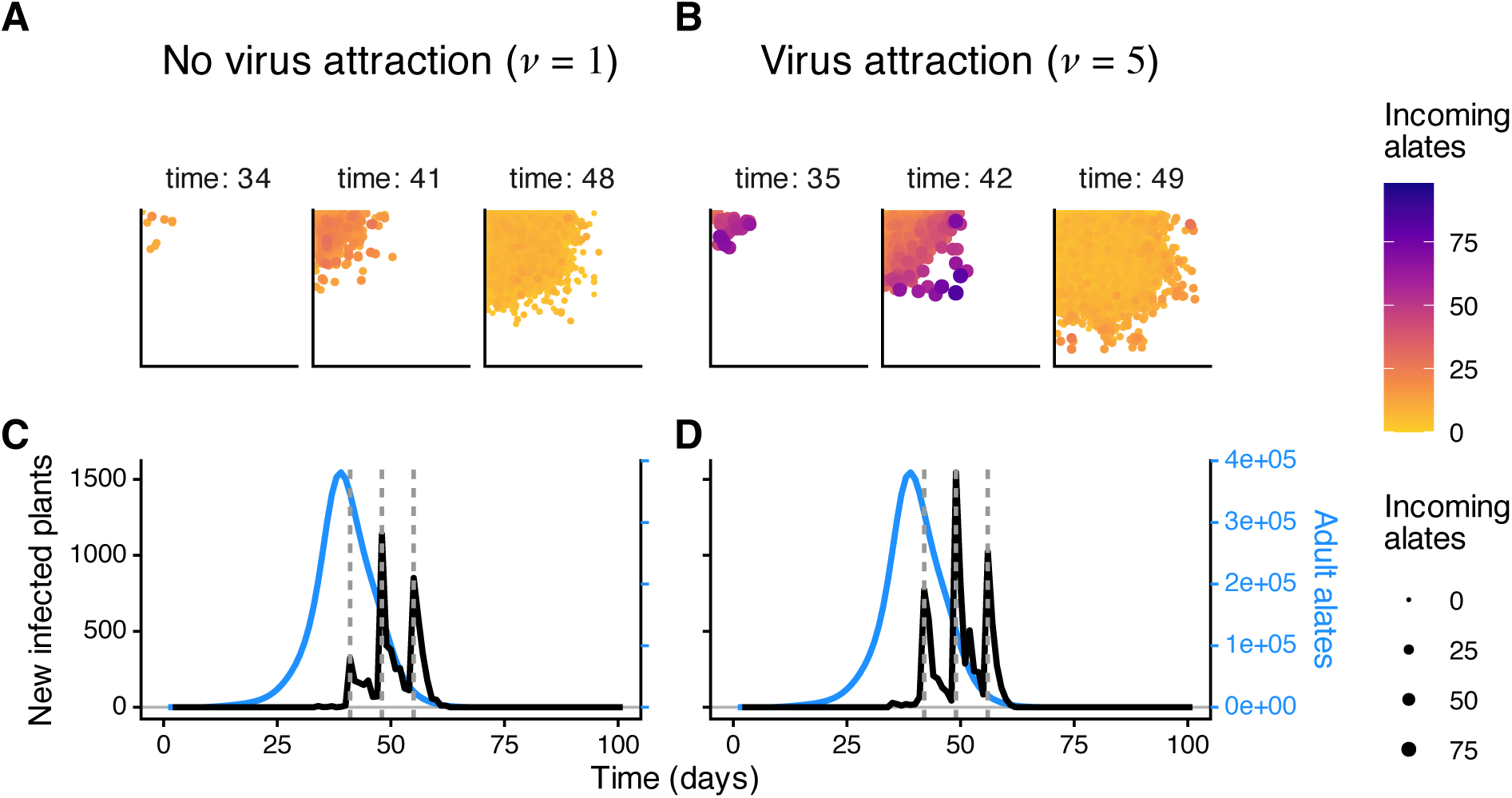
Virus attraction accelerates the traveling wave of virus infection in large 100 × 100 landscapes. (A,B) Locations and numbers of incoming alates for virus-infected plants when virus attraction is (A) absent (*ν* = 1) or (B) present (*ν* = 5). The 3 chosen time points are those preceding the largest peaks in new infected plants by 7 days (the incubation period for the virus in our model) for each set of simulations. These days thus represent the alate dispersal events that caused these new-infection peaks. Each point indicates a virus-infected plant, and both color and size of points indicate the number of incoming alates visiting and probing (but not necessarily staying to feed at) plants. Panels separate time points, as indicated above each panel. (C,D) Line plots show the (black lines) numbers of newly infected plants and (blue lines) adult alate densities through time, when virus attraction is (C) absent or (D) present. The vertical dashed lines indicate the peaks in new infections 7 days following each time in panels A and B. For all panels, *Pseudomonas* is absent from these simulations to exclude its effects, and parasitoid responsiveness was set to a moderate value (*ζ* = 0.5) because it was not of interest here. Loading probabilities (*δ_a_*, *δ_p_*) were set to 1 to maximize virus prevalence, setting up a scenario where virus attraction is typically thought to decrease virus transmission. Other parameters and starting conditions are the same as for Figure 4.

**Figure S8:**
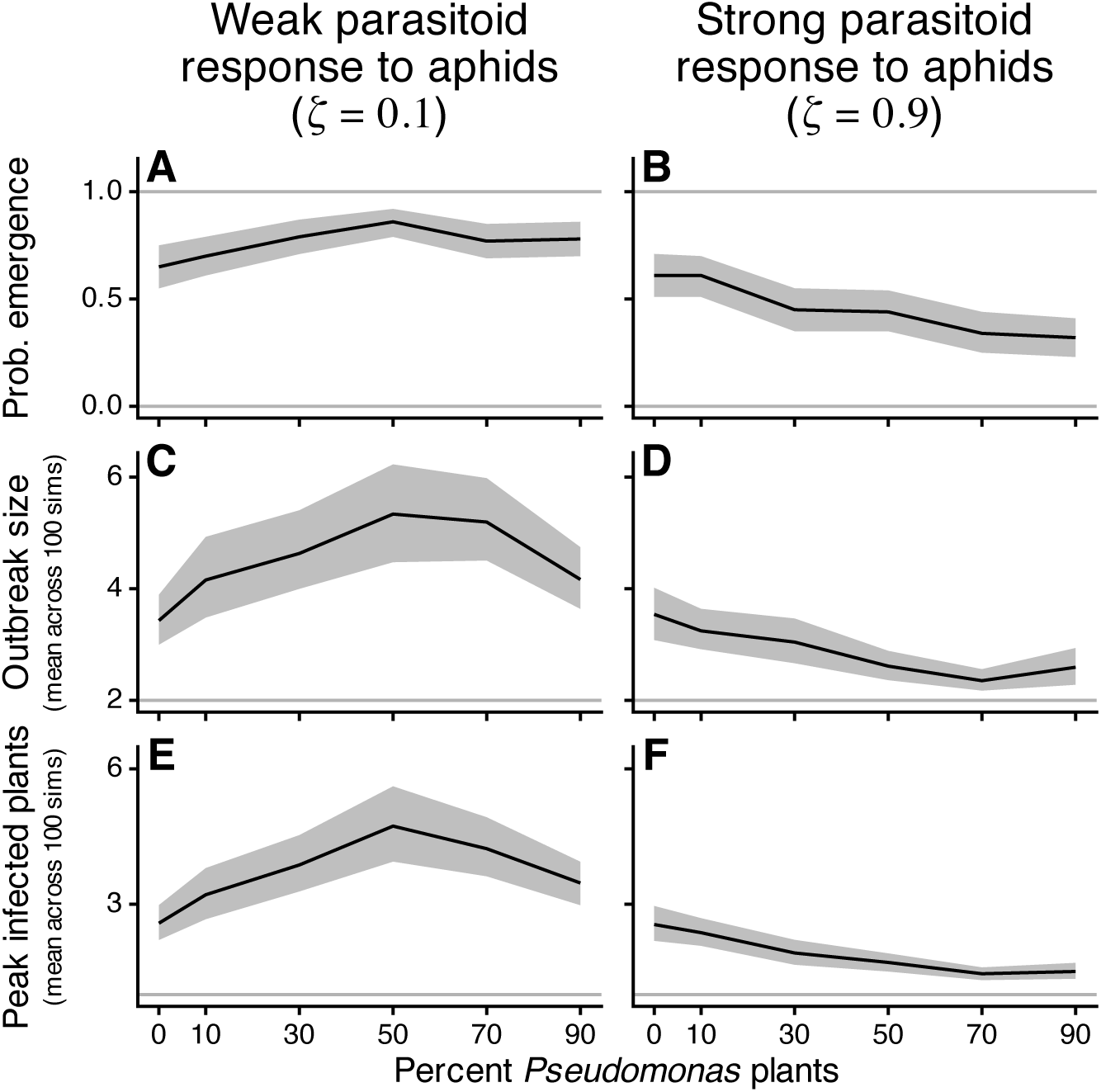
In larger 100 × 100 landscapes of pea plant plants where we simulate conditions where outbreaks do not always occur, the percent of plants with *Pseudomonas* affects virus outbreaks. Results are shown for (A,C,E) weak (*ζ* = 0.1) and (B,D,F) strong (*ζ* = 0.9) parasitoid responsiveness to aphid densities. Line plots show the (A,B) probability of emergence, (C,D) mean outbreak size, and (E,F) mean peak infected plants across 100 simulations versus percent *Pseudomonas* plants. Shaded ribbons indicate 95% confidence intervals. Parameters and starting conditions are the same as for Figure 4, except that *δ_a_* = *δ_p_* = 0.05.

**Figure S9:**
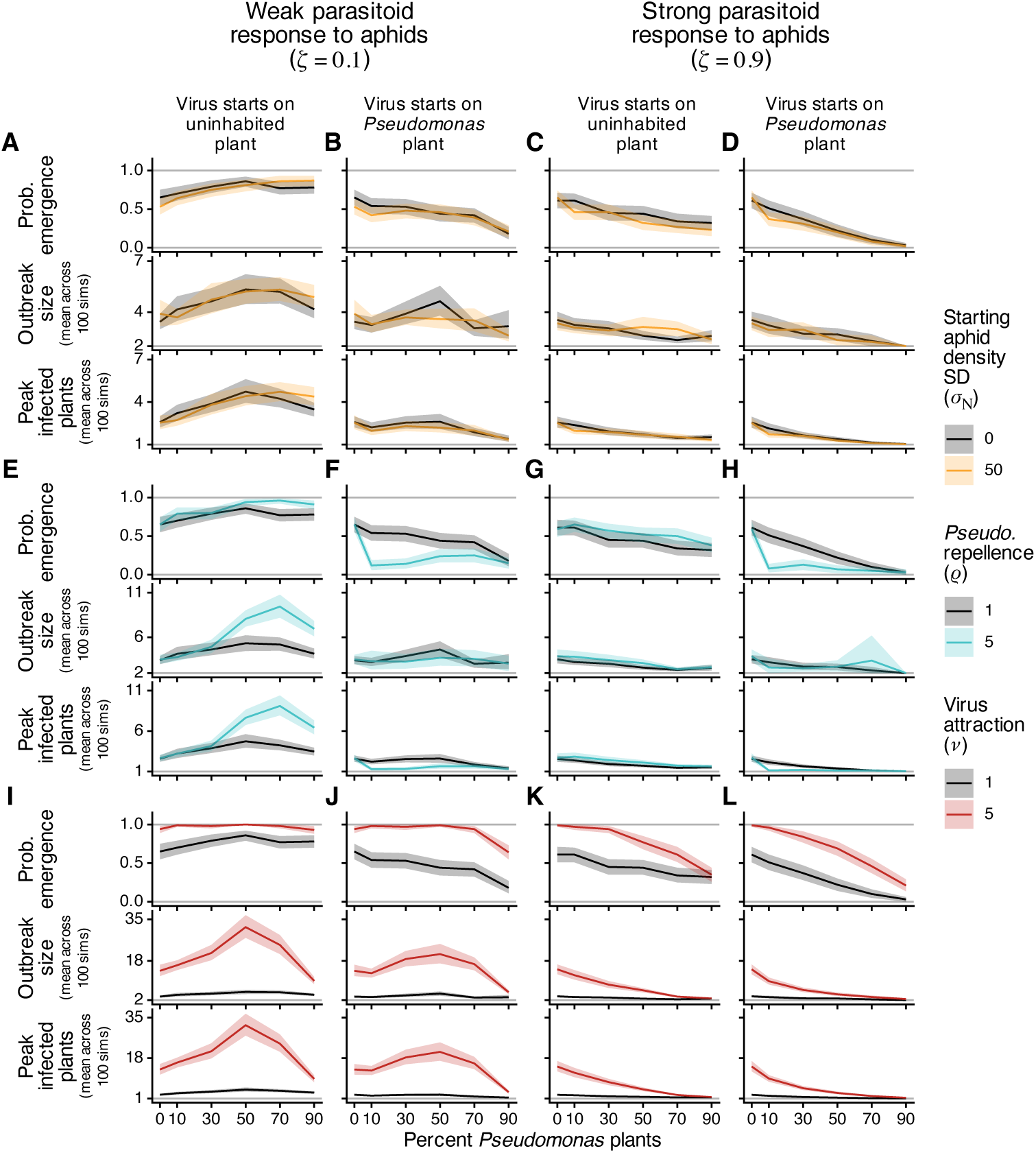
In larger 100 × 100 landscapes of pea plant plants with reduced loading probabilities (*δ_a_* = *δ_p_* = 0.05 instead of 0.5) that cause outbreaks to not always occur, the effect of *Pseudomonas* on virus outbreak sizes can be altered by factors affecting the spatial distribution of insects and viruses. Results are shown for (A,B,E,F,I,J) weak (*ζ* = 0.1) and (C,D,G,H,K,L) strong (*ζ* = 0.9) parasitoid responsiveness to aphid densities. Line plots show the (top panel) probability of emergence, (middle) mean outbreak size, and (bottom) mean peak infected plants across 100 simulations versus percent *Pseudomonas* plants. In all panels, the black line is results from the baseline simulations where attractiveness to alates and initial aphid densities are uniform across plants. Each colored line is from simulations where we change one component of the model: (A–D) there is variation in starting aphid densities (*σ_N_* = 50), (E–H) *Pseudomonas* repels alates (*ρ* = 5), or (I–L) virus infection attracts alates (*ν* = 5). Note the differences in scales of the y-axes among panels for mean outbreak size and peak infected plants. Parameters and starting conditions are the same as for Figure S8, except for loading probabilities and those indicated for the colored lines.

**Figure S10:**
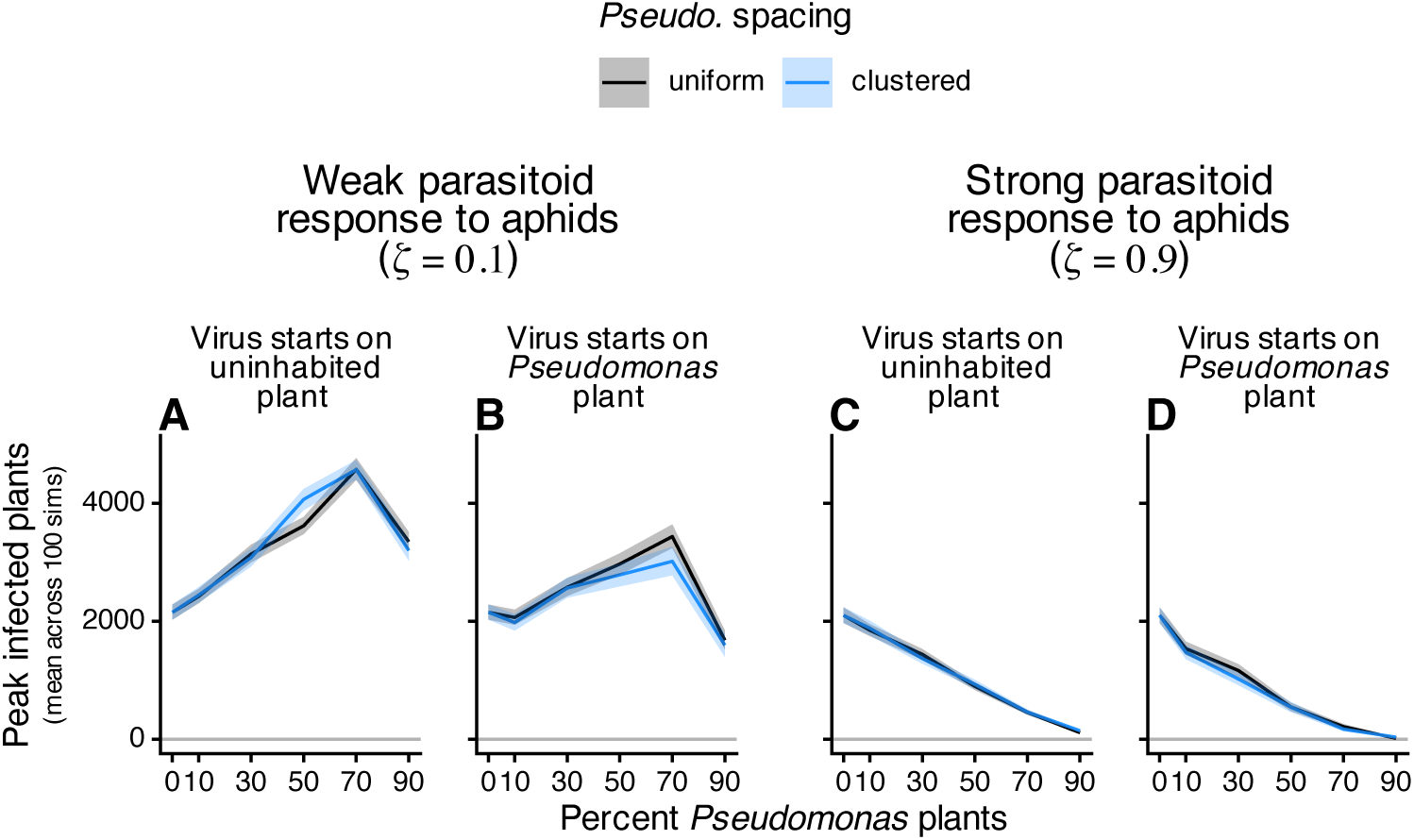
In larger 100 × 100 landscapes of pea plant plants, the effect of *Pseudomonas* on outbreaks is not significantly altered by clustering of *Pseudomonas* locations. Results are shown for (A,B) weak (*ζ* = 0.1) and (C,D) strong (*ζ* = 0.9) parasitoid responsiveness to aphid densities, and for whether the virus starts on an (A,C) uninhabited or (B,D) *Pseudomonas*-inhabited plant. Line plots show the mean peak infected plants across 100 simulations, versus percent *Pseudomonas* plants. In all panels, the black line is baseline simulations where *Pseudomonas* locations are uniformly located on the landscape. Blue lines are from simulations where *Pseudomonas*-inhabited plant locations are spatially clustered instead of uniformly distributed. See Section S3 for how clustering was simulated. Parameters and starting conditions are the same as for Figure 4, except for those indicated.

**Figure S11:**
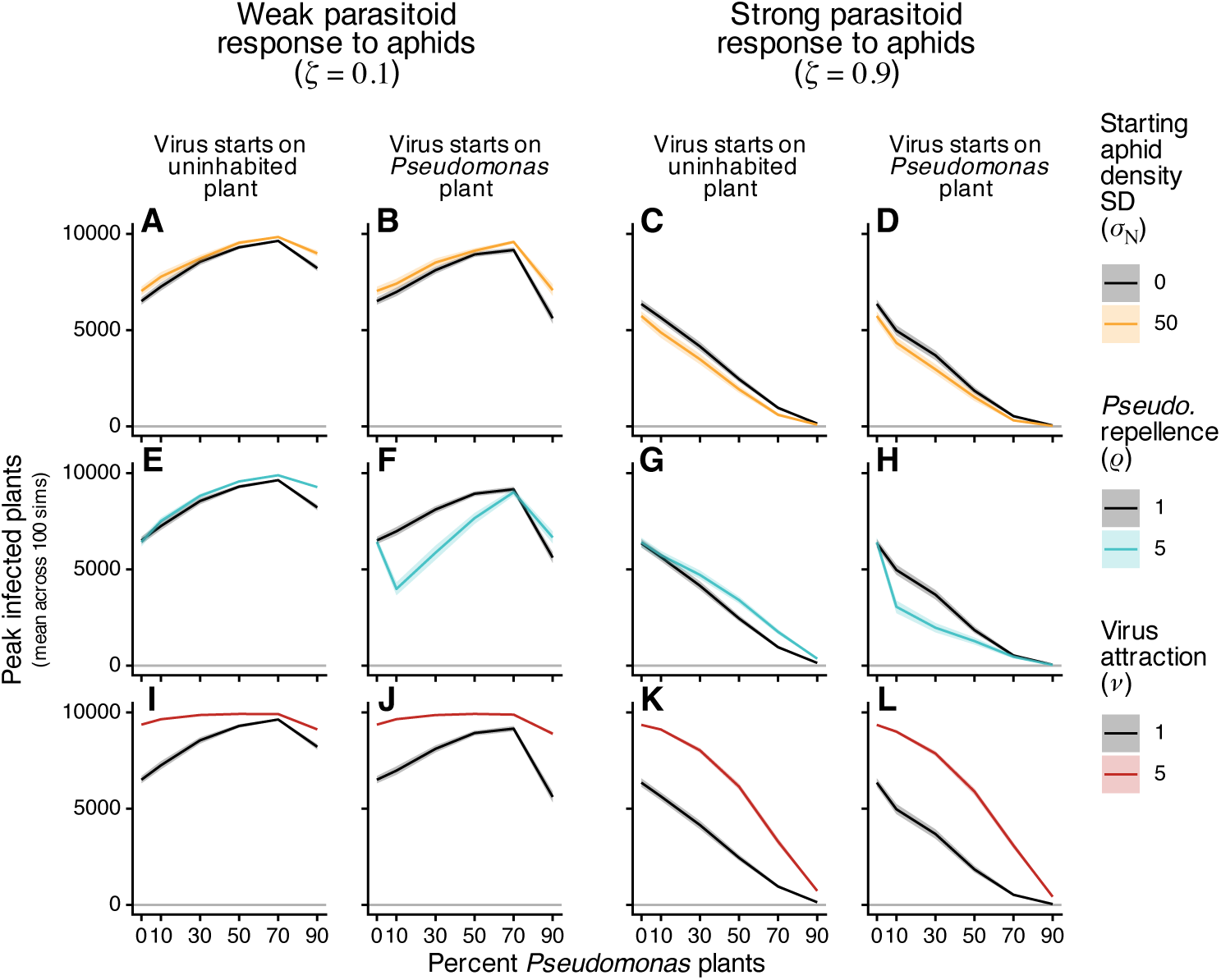
In larger 100 × 100 landscapes of pea plant plants where we start the virus on an interior plant (*x* = 50, *y* = 50), the effect of *Pseudomonas* on virus outbreak sizes can be altered by factors affecting the spatial distribution of insects and viruses. Results are shown for (A,B,E,F,I,J) weak (*ζ* = 0.1) and (C,D,G,H,K,L) strong (*ζ* = 0.9) parasitoid responsiveness to aphid densities, and for whether the virus starts on an (A,C,E,G,I,K) uninhabited or (B,D,F,H,J,L) *Pseudomonas*-inhabited plant. Line plots show the mean peak infected plants across 100 simulations versus percent *Pseudomonas* plants. In all panels, the black line is results from the baseline simulations where attractiveness to alates and initial aphid densities are uniform across plants. Each colored line is from simulations where we change one component of the model: (A–D) there is variation in starting aphid densities (*σ_N_* = 50), (E–H) *Pseudomonas* repels alates (*ρ* = 5), or (I–L) virus infection attracts alates (*ν* = 5). Probability of emergence and mean outbreak size are not shown because outbreaks nearly always occurred in these simulations. Parameters and starting conditions are the same as for Figure 4, except for viruses starting on interior plants and others as indicated for the colored lines.

**Figure S12:**
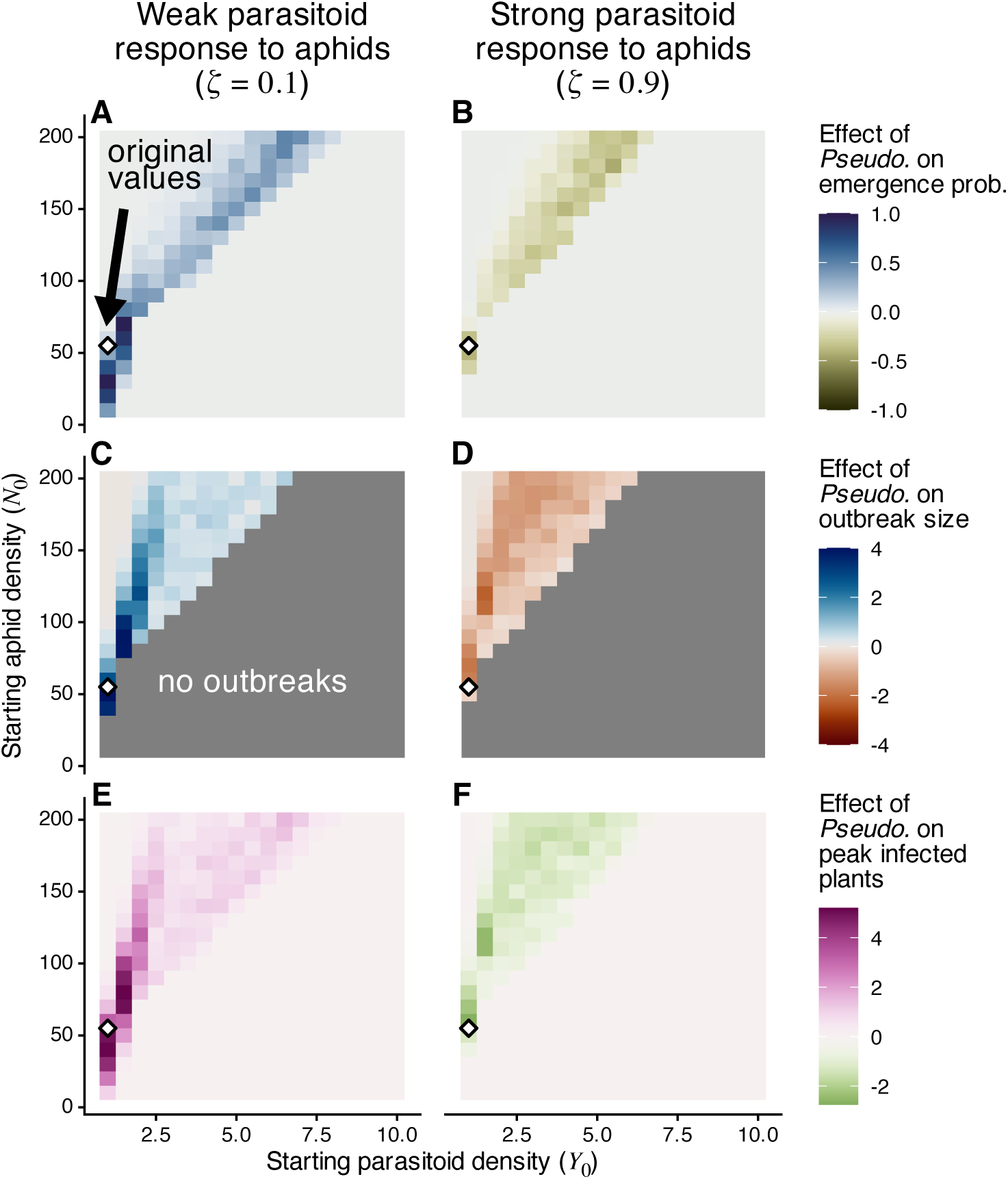
Heatmaps showing how *Pseudomonas* presence affects (A,B) probability of emergence (i.e., an outbreak occurring) and (C,D) mean outbreak size (when an outbreak occurred), and (E,F) mean peak infected plants across starting densities of parasitoids (*Y*_0_) and aphids (*N*_0_). Outbreaks are defined as at least one new plant being infected, and the effect of *Pseudomonas* is measured by each outcome with *Pseudomonas* minus that without. Results are shown for (A,C) weak (*ζ* = 0.1) and (B,D) strong (*ζ* = 0.9) parasitoid responsiveness to aphid densities. White diamonds indicate the values used for the original simulations shown in Figure 2. (C,D) Combinations that resulted in no outbreaks either with or without *Pseudomonas* are gray.

**Figure S13:**
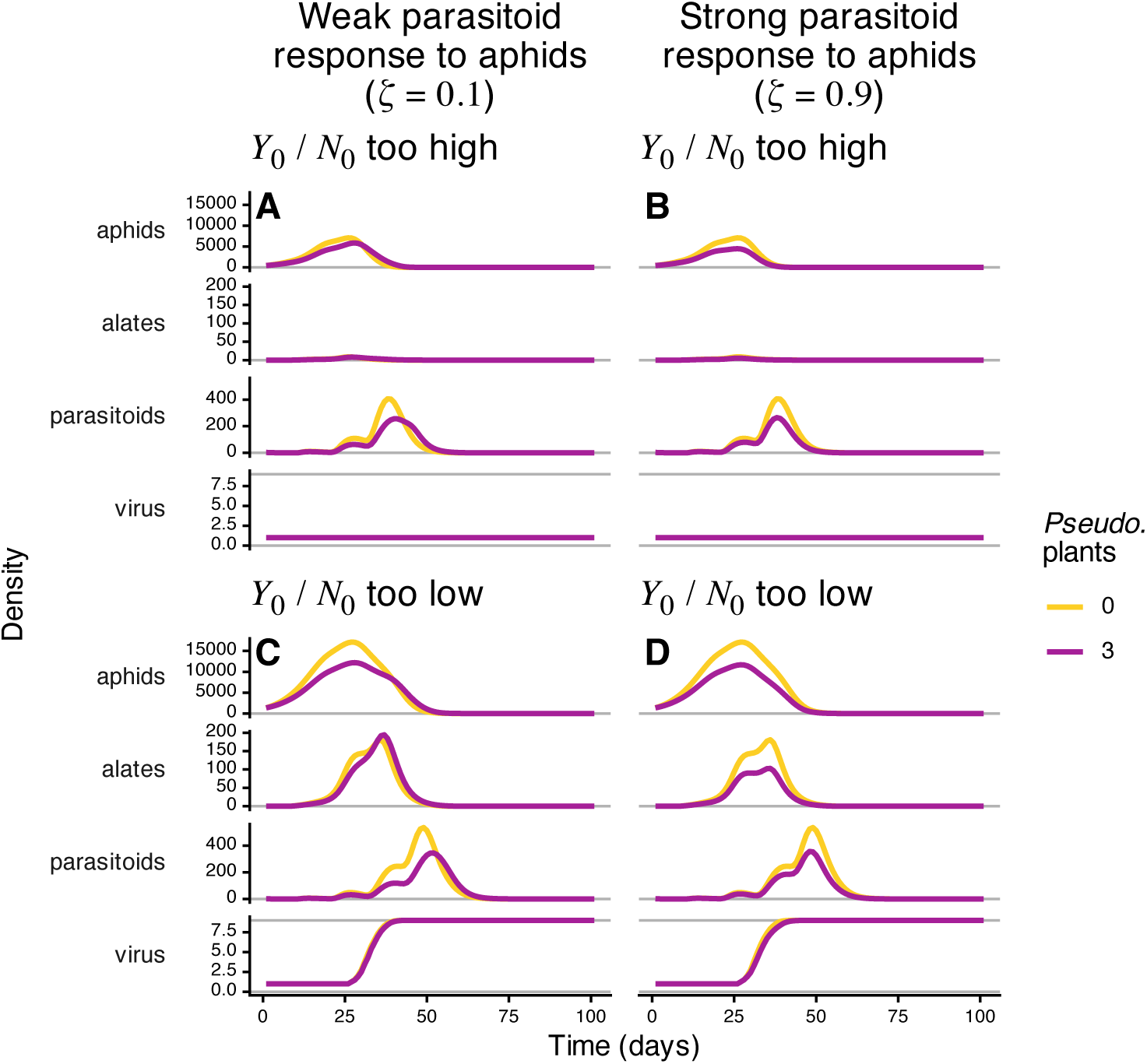
For simulations where starting parasitoid densities relative to starting aphid densities are either (A,B) too high or (C,D) too low, time series of total landscape-wide density for total aphids, adult alates, adult parasitoids (female only), and plants infectious with the virus. Results are shown for parasitoid responsiveness to aphid densities being (A,C) weak (*ζ* = 0.1) or (B,D) strong (*ζ* = 0.9). Gold lines indicate no *Pseudomonas* on the landscape, and magenta lines indicate 3 *Pseudomonas*-inhabited plants. Parameters and starting conditions are the same as for Figure 2, except for the following: When above the threshold, *Y*_0_ = 3 and *N*_0_ = 50, and when below the threshold, *Y*_0_ = 1 and *N*_0_ = 150. Starting densities do not differ between panels of the same row.

**Figure S14:**
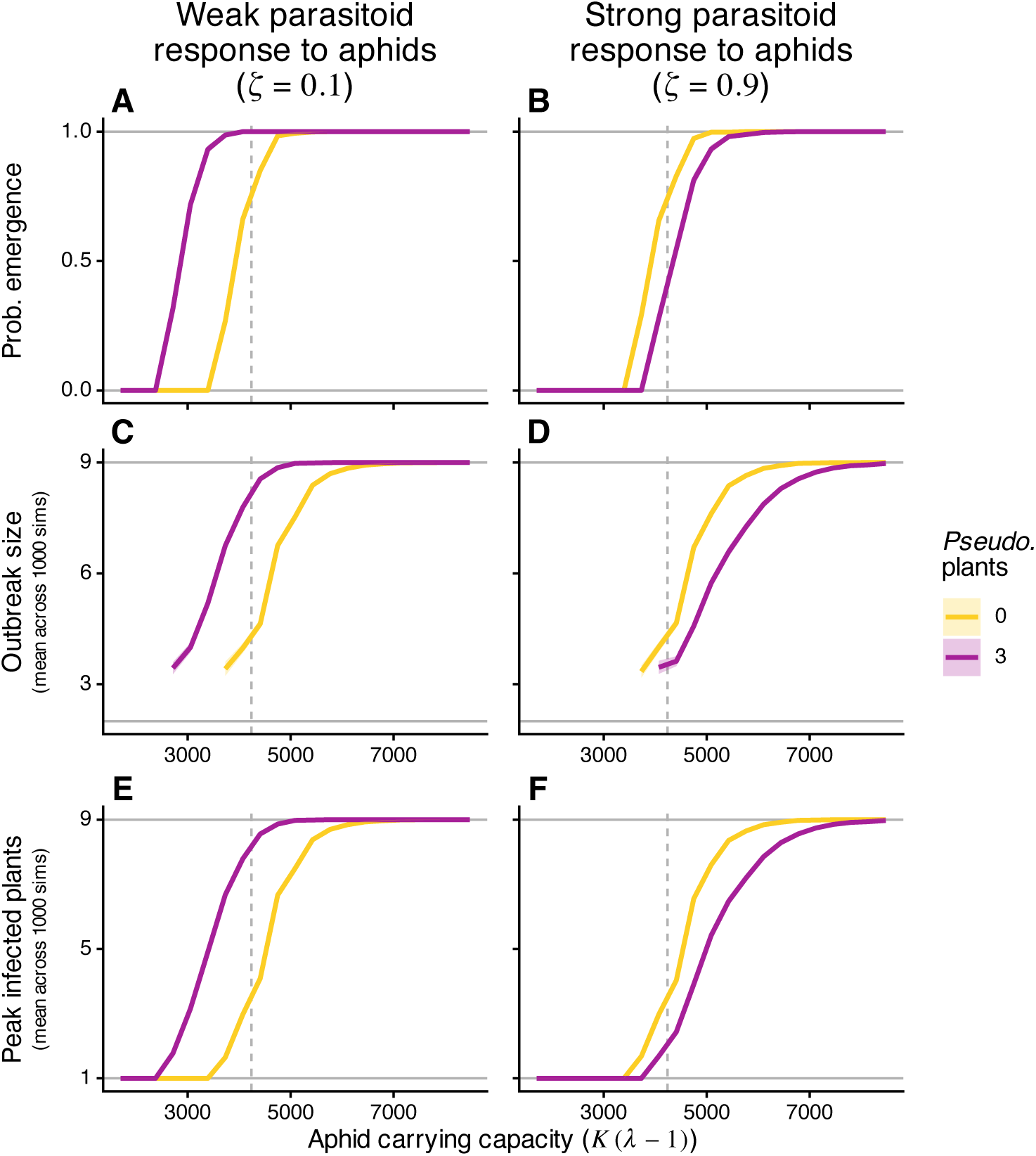
Line graphs showing the (A,B) probability of emergence and (C,D) mean outbreak size, and (E,F) mean peak infected plants across values of aphid carrying capacity (*K* (*λ* − 1)), for parasitoids responding (A,C,E) weakly (*ζ* = 0.1) or (B,D,F) strongly (*ζ* = 0.9) to aphid densities. Lines are shown for landscapes (magenta) with and (gold) without *Pseudomonas*-inhabited plants. Vertical dashed lines indicate the carrying capacity values used for the original simulations shown in Figure 2. In panels C and D, lines are truncated for carrying capacities where outbreaks never occur. Shaded ribbons indicate 95% confidence intervals.

**Figure S15:**
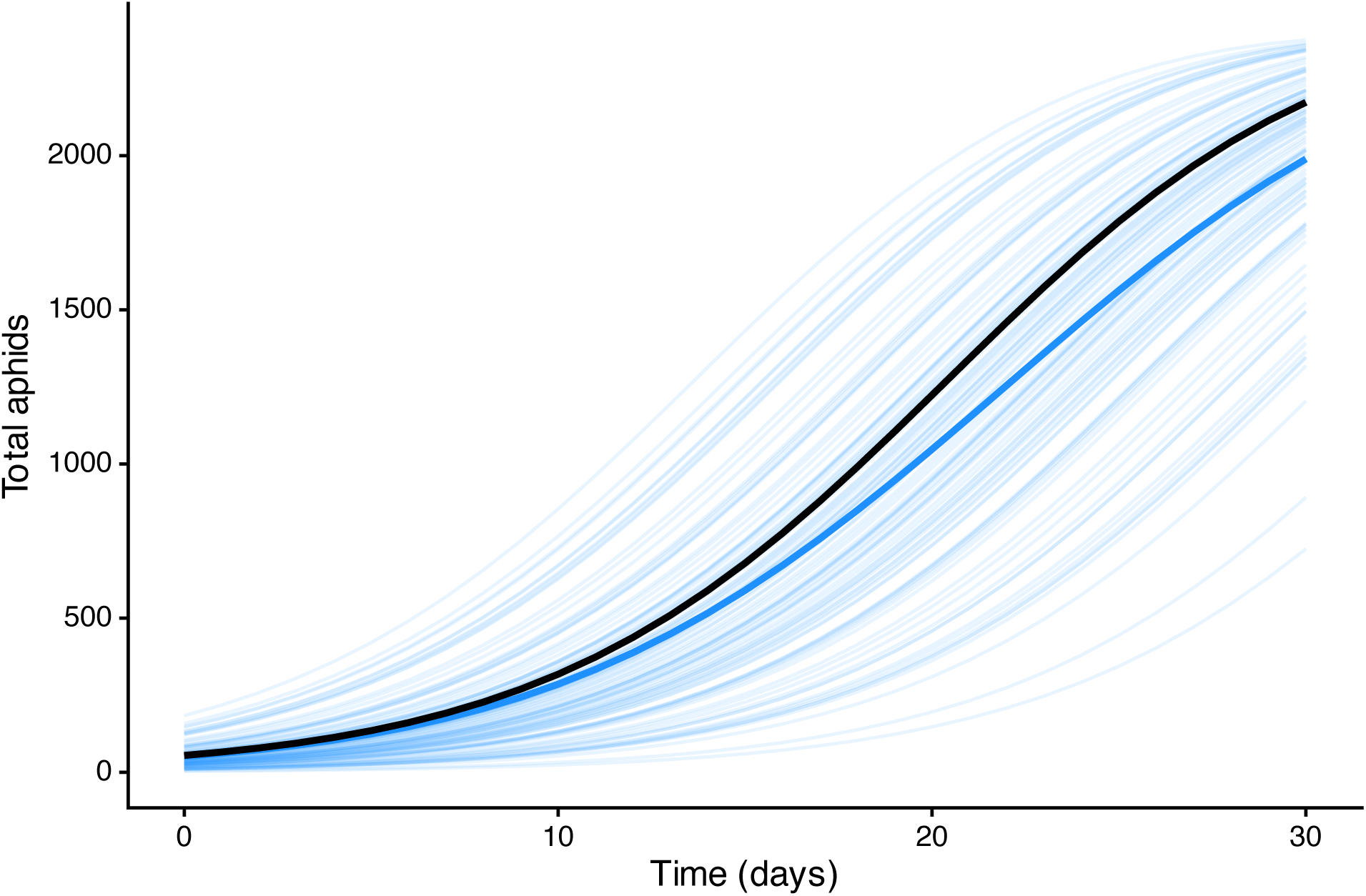
Time series showing the effect of variability in initial aphid densities on total aphid densities for the beginning of a growing season. Thin, blue lines are total aphid densities for the 100 individual plants for which starting aphid densities were generated from a lognormal distribution with a mean of 55 and standard deviation of 50 (see “Methods” for details). The thick blue line is the mean of these plants through time. The thick black line is aphid densities for a plant with a starting aphid density of 55. These simulations focused only on aphid populations, so they did not include parasitoids, *Pseudomonas*, or virus transmission. Other than those indicated, starting conditions and parameter values matched those from Figure 2.

## Notes

### Competing Interest Statement

The authors have declared no competing interest.

### Summary of Updates

We streamlined the narrative, with many of the previous figures and results (and some methods) going into the supplement. We also improved the discussion for flow and content.

